# Mechanistic insights into the success of xenobiotic degraders resolved from metagenomes of microbial enrichment cultures

**DOI:** 10.1101/2021.03.03.433815

**Authors:** Junhui Li, Chongjian Jia, Qihong Lu, Bruce A. Hungate, Paul Dijkstra, Shanquan Wang, Cuiyu Wu, Shaohua Chen, Deqiang Li, Hojae Shim

**Author notes:** Corresponding authors (J. Li), (H. Shim).

## Abstract

Even though microbial communities can be more effective at degrading xenobiotics than cultured micro-organisms, yet little is known about the microbial strategies that underpin xenobiotic biodegradation by microbial communities. Here, we employ metagenomic community sequencing to explore the mechanisms that drive the development of 49 xenobiotic-degrading microbial communities, which were enriched from 7 contaminated soils or sediments with a range of xenobiotic compounds. We show that multiple microbial strategies likely drive the development of xenobiotic degrading communities, notably (i) presence of genes encoding catabolic enzymes to degrade xenobiotics; (ii) presence of genes encoding efflux pumps; (iii) auxiliary catabolic genes on plasmids; and (iv) positive interactions dominate microbial communities with efficient degradation. Overall, the integrated analyses of microbial ecological strategies advance our understanding of microbial processes driving the biodegradation of xenobiotics and promote the design of bioremediation systems.

## 1. Introduction

Environmental contamination by xenobiotics (e.g., aromatic hydrocarbons, chlorinated solvents) has been a serious global issue for over half a century. Microorganisms play significant roles in the cleanup of these contaminants (Singh and Ward 2004). Extensive research on isolated pure cultures has provided insights into microbial metabolism by characterizing genes encoding specialized enzymes and biodegradation pathways for contaminants (de Lima-Morales et al. 2016, Suttinun et al. 2013, Zhou et al. 2018). Microbial communities often break down xenobiotics more efficiently than pure cultures, likely due to synergistic cooperation which increases metabolic capabilities, more efficient resource utilization, elevated ecological stability, and enhanced survivability (Ben Said and Or 2017, Cerqueira et al. 2011, Faust 2019, Garcia 2016, Hays et al. 2015, Kang et al. 2020, Lindemann et al. 2016, Xu et al. 2019), and thus communities are well suited for real-world applications (Hays et al. 2015). Advances in high-throughput sequencing have provided a comprehensive survey of the diversity and composition of microbial communities, yet little is known regarding the potential mechanisms underlying the biodegradation process, hindering our ability to harness microbiomes to detoxify environmental contaminants.

Microbial ecologists seek to understand the microbial mechanisms that contribute to biogeochemical cycles, and trait-based approaches have expanded our understanding of ecological patterns and ecosystem functioning (Krause et al. 2014, Lajoie and Kembel 2019, Li et al. 2019, Malik et al. 2020, Martiny et al. 2015, Sorensen et al. 2019). Metabolic trait dissimilarities reflect strategies for acquiring resources among species and many microbial traits seem to be phylogenetically conserved (Martiny et al. 2015), including carbon (Berlemont and Martiny 2015, Martiny et al. 2015) and nitrogen (Isobe et al. 2019, Martiny et al. 2015) cycling. Genes encoding catabolic enzymes contribute to the biodegradation of xenobiotics (van der Meer et al. 1992). Besides catabolic enzymes, other microbial ecological strategies that govern xenobiotics breakdown remain largely unknown. Bacteria can acquire genes via plasmid-mediated horizontal gene transfer (HGT) in microbial communities to adapt to environmental changes (Hall et al. 2016) and plasmid-mediated HGT plays a vital role in petroleum hydrocarbon biodegradation (Shahi et al. 2017) as genes encoding the degradation of organic contaminants are often located on plasmids (Garbisu et al. 2017). On the other hand, stress tolerance has been proposed as a microbial life strategy in the study of soil carbon cycling (Krause et al. 2014, Malik et al. 2020). Microorganisms exposed to organic solvents have developed adaptation mechanisms to overcome the solvent stress (Kusumawardhani et al. 2018, Rojo 2016). For instance, efflux pumps of multiple strains of *Pseudomonas putida* confer resistance to aromatic solvents (Basu et al. 2006, Inoue and Horikoshi 1989, Molina-Santiago et al. 2017, Sol Cuenca et al. 2015), among which *P. putida* CSV86 demonstrated preferential utilization of aromatics over glucose (Basu et al. 2006). During long-term exposure to xenobiotics, microbial communities are subject to evolution and adaption that can result in the acquisition of novel catabolic enzymes and efflux pumps via hosting of plasmids, and thus could gain the ability to utilize xenobiotics as growth substrates (Poursat et al. 2019). In nature, microbial competition for resources is common, but complex substrates (e.g., cellulose, xylan and lignin) promote synergistic interactions (Deng and Wang 2017). This has been observed in the biodegradation of xenobiotics as well (Leewis et al. 2016). Information about these microbial traits could be particularly useful in linking microbial processes to the biodegradation of xenobiotics. Individual microbial traits, particularly catabolic traits, have been often studied in xenobiotic biodegradation, however the integration of multiple microbial traits of the microbial community assembly has rarely been considered.

Previously we investigated the community composition of 49 enriched microbial communities, grown for one year with a range of contaminants and diverse origin using 16S rRNA gene amplicon sequencing, and efficient aromatic biodegradation was achieved (Li et al. 2020). We found that the genera *Pseudomonas* and *Rhodococcus* were the prevalent aromatic degraders (Li et al. 2020). We applied whole genome shotgun sequencing to identify the potential mechanisms driving the success of major xenobiotics degraders. We hypothesized four mechanisms (Fig. 1) responsible for the development of enriched degraders, i.e., 1) metabolism via catabolic enzymes; 2) gene acquisition via plasmid-mediated HGT; 3) solvent stress tolerance through efflux pumps; and 4) synergistic interactions among enriched community members. Integrated analyses at the community level, genus level, genome level (i.e., genome bins), and replicon level (i.e., plasmid and chromosome) were performed to test the hypotheses.

**Fig. 1.**
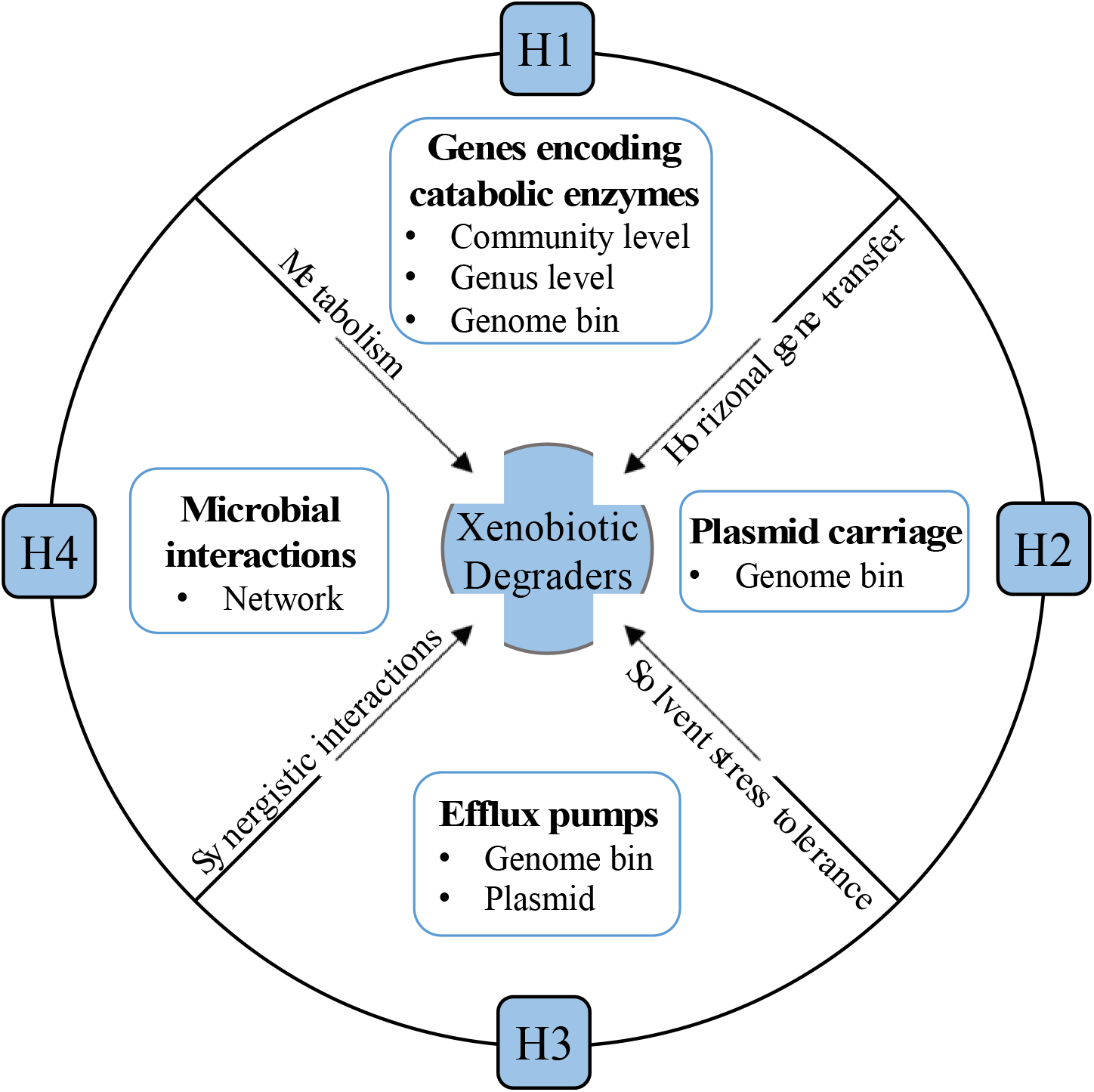
Hypothesized (H1-4) mechanisms driving the success of aerobic cultivatable xenobiotic degraders over one-year incubation.

## 2. Materials and Methods

### 2.1. Laboratory incubation, DNA extraction, and sequencing

Forty-nine xenobiotic degrading microbial communities enriched from seven contaminated soil or sediment samples, together with 12 mixed cultures (see below), were collected for metagenomic sequencing. (Table S1). Details of laboratory incubation (enrichment of microbial communities and DNA extraction) are described elsewhere (Li et al. 2020). In short, soils and sediments were firstly enriched in nutrient broth, and then enriched in mineral salts medium supplying different xenobiotic contaminants as substrates. The microbial communities were sub-cultured with xenobiotics over a year on a weekly basis (Table S1). DNA was extracted from each enrichment culture at the end of the respective incubation period using the MP FastDNA™ SPIN Kit. The DNA quality and quantity were estimated using the Implen NanoDrop (Munich, Germany). Libraries were prepared using the Nextera DNA Flex Library Preparation kit (Illumina Inc., San Diego, CA, USA). Next, the pooled libraries were sequenced on the Illumina NovaSeq 6000 platform (Illumina Inc., SanDiego, CA, USA) with 150 bp paired-end reads. Sequencing service was provided by Berry Genomics Co. Ltd. One sample (QLP.T2.1) was sequenced twice due to the low sequencing depth of the first sequencing, and data from two sequencing runs were combined.

### 2.2. Bioinformatic analyses

Raw reads were trimmed using TrimGalore (Krueger 2015) with default settings, then reads were aligned to the human reference genome hg38 using Bowtie2 (Langmead and Salzberg 2012) to filter out human sequences. Finally, a total of 1.98 billion reads were obtained, corresponding to an average of 32.5 million paired-end reads per sample. All metagenomes from the same sample source were co-assembled separately using Megahit (Li et al. 2015). On average, 55,656 contigs with a minimum length of 500 bp were obtained across the co-assemblies, with N50 of 11,071. Clean sequences were then mapped to the co-assembled contigs from the respective sample source using BWA (Li and Durbin 2009) and SAMtools (Li et al. 2009). Contigs were taxonomically annotated using Kaiju (Menzel et al. 2016) [−a greedy −e 3 −E 0.00001]. For the catabolic genes and efflux pumps, we predicted the open reading frames (ORFs) of the contigs using Prodigal (Hyatt et al. 2010) [metagenomic mode]. Subsequently, predicted ORFs were searched using hmmscan in HMMER V3.2.1 (Finn et al. 2011) [E-value < 1e-15 and covered fraction of HMM > 0.5] against the targeted KO families that are mapped to KEGG metabolic pathways of xenobiotics (map00361 for benzene; map00623 for toluene; map00642 for ethylbenzene; map00622 for xylenes; map00625 for chlorinated solvents) from KOfam database (Aramaki et al. 2020), together with three self-constructed hmm files (DMF.hmm for *N*,*N*-dimethylformamide (DMF), EaCoMT.hmm and vcrC.hmm for chlorinated solvents), which were built using the ‘hmmbuild’ command based on the multiple protein sequence alignments [by Clustal Omega V1.2.4 (Sievers and Higgins 2014)]. The constructed HMM files and their seed protein sequences are available at https://bitbucket.org/junhuilinau/hmm/src/master/. Next, we used a subset of the Resfams (Gibson et al. 2015) families to identify the efflux pumps with hmmscan (Finn et al. 2011) [--cut_ga]. The functional abundances were normalized by the total annotated sequences, including bacteria (99.98%), viruses (0.012%), fungi (0.0070%), and archaea (0.00087%). One sample (T6 from petrochemical complex, Table S1) was excluded from community level analyses due to preparation error.

Further, each metagenome was assembled individually using Megahit (Li et al. 2015), and the resulting contigs, together with co-assemblies, were then binned with metaBAT2 (Kang et al. 2019), MaxBin2 (Wu et al. 2016), and CONCOCT (Alneberg et al. 2014) and refined with the metaWRAP (Uritskiy et al. 2018) pipeline, and bins < 70% completeness or > 5% contamination according to CheckM (Parks et al. 2015) were removed and dereplicated using dRep (Olm et al. 2017), resulting in a total of 201 metagenomic assembled genomes (MAGs) with the average nucleotide identity < 99.8% except four major genera [i.e., keeping all bins unless with identical 40 marker genes (Wu et al. 2013)]. The 201 obtained MAGs were estimated have on average 93.3% completeness, 1.2% contamination, G+C content of 0.640 ranging between 0.376 and 0.714, and N50 of 53.3 kb. The functional annotation (i.e., catabolic genes and efflux pumps) of MAGs was determined in the same way as for contigs. The taxonomic annotations are from the phylogenetic classification based on 40 single copy universal marker genes as indicated below. Plasmid carriage on MAGs was predicted using Platon V1.2.0 (Schwengers et al. 2020).

### 2.3. Phylogenetic visualization

A phylogenetic tree including representative species from all 199 genera that harbor catabolic genes was obtained from NCBI common tree. The phylogenetic trees of MAGs were constructed using iqtree (Minh et al. 2020) under the MFP model with 40 single copy universal marker genes (Wu et al. 2013), which were extracted from the predicted ORFs in the MAGs using fetchMGs v1.2 (Mende et al. 2013), aligned using Clustalo (Sievers et al. 2011), and trimmed using trimAl v1.2 (Capella-Gutiérrez et al. 2009) [-gt 0.5]. iTOL (Letunic and Bork 2016) was used for visualizing the phylogenetic trees.

### 2.4. Correlation network

Owing to the distinct beta diversity between treatments in the presence and absence of BTEX (benzene, toluene, ethylbenzene, *o*-xylene, *m*-xylene, and *p*-xylene), two microbial co-occurrence networks were constructed using taxa that are present in at least half of the metagenomes (presence 900 taxa; absence 1,047 taxa), accounting for the average of around 99% of the community composition, irrespective of the presence or absence of BTEX. The conservative cut-off of Spearman ρ > 0.80 and *p* < 0.01 (multiple testing adjustment using Benjamini-Hochberg correction) were used to generate statistically robust correlations and control the false-positive rate. Gephi (Bastian et al. 2009) was used to visualize the networks, and modularity was estimated with the modularity function using the Louvain clustering algorithm (Blondel et al. 2008).

### 2.5. Data availability

Metagenomic sequence data were deposited in the NCBI sequence read archive under accession No. PRJNA660264. All other data that support the findings of this study are available from the corresponding author upon reasonable request.

### 2.6. Statistical analyses

All the statistical analyses were performed with R (https://www.r-project.org/). The permutational multivariate analysis of variance (PERMANOVA) was performed with Bray-Curtis distance and binary Jaccard distance using adonis2 function in Vegan package (Oksanen et al. 2013). The nonmetric multidimensional scaling (NMDS) coordination with abundance‐based Bray–Curtis and presence/absence‐based binary Jaccard distances (permutations = 9,999) was generated to visualize the differences in microbial community and catabolic genes. All the figures were generated in the ggplot2 package aside from the phylogenetic tree and network.

## 3. Results

### 3.1. Taxonomic and functional compositions of microbial communities predict substrates, while metagenomic community gene diversity predicts inoculum sources

The microbial communities are composed of bacteria, fungi, archaea and viruses while dominated by bacteria (> 99.9%), and all the microbial taxa were included in analyses unless otherwise stated. As demonstrated in the supplementary, the metagenomic genera generally display a high degree of consistency to 16S rRNA gene amplicon sequencing in the enriched microbial communities (Fig. S1, Fig. S2). Similar to 16S amplicon sequences (R^2^ = 0.817, *p* < 1e-4, Bray-Curtis distance, Fig. 2a), the metagenomic microbial community composition differs significantly among treatments with different xenobiotics based on genus-level taxonomy and Bray-Curtis beta diversity (R^2^ = 0.816, *p* < 1e-4, Fig. 2b). Likewise, the enriched microbial communities from different treatments vary greatly in community catabolic gene abundances (R^2^ = 0.405, *p* < 1e-4, Bray-Curtis distance, Fig. 2c). Consistently, the treatments in the presence and absence of BTEX are differentiable in the genus compositions and community catabolic potential (Fig. 2a-c). In addition to the community composition, dominant taxa varied substantially between treatments in the presence and absence of BTEX, i.e., *Pseudomonas* (average 55.8%) and *Rhodococcus* (23.5%) were the dominant genera in the treatments in the presence of BTEX, while *Paracoccus* (70.2%) and *Hyphomicrobium* (10.9%) were the dominant genera in the treatments in the absence of BTEX (Fig. S2). The 16S amplicon analysis of dominant genera supported this finding (Fig. S2). Moreover, the inoculum source also causes a significant difference in the microbial community compositions, albeit small proportions of explained variation (16S amplicon: R^2^ = 0.062, *p* = 7e-03, Bray-Curtis distance, Fig. 2a; shotgun metagenome: R^2^ = 0.055, *p* = 0.028, Bray-Curtis distance, Fig. 2b), and the community catabolic potential (R^2^ = 0.198, *p* = 0.0012, Bray-Curtis distance, Fig. 2c). Likewise, although the presence or absence of BTEX dominates in shaping the microbial enrichment cultures, inoculum sources and other treatments significantly affected the community composition (Fig. S2).

As compared to the analyses based on Bray-Curtis beta diversity (0.055 – 0.198, *p* ≤ 0.028, Fig. 2a-c), the inoculum source explains a greater proportion of variation in the binary Jaccard beta diversity (0.213 – 0.790, *p* < 1e-4, Fig. 2d-f). Moreover, the inoculum source exerted stronger influence on the binary Jaccard beta diversity of metagenome at the genus level (R^2^ = 0.457, *p* < 1e-4, Fig. 2e) than substrates (R^2^ = 0.244, *p* < 1e-4, Fig. 2e). Notably, the inoculum source explains 79% of the variation in binary Jaccard beta diversity of community genes (*p* < 1e-4, Fig. 2f). Overall, our results suggest that taxonomic and functional Bray-Curtis beta diversities predict the presence or absence of the added contaminants, while binary Jaccard beta diversity (i.e., richness) of catabolic genes separates samples according to their inoculum sources. Thus, taxonomic and functional beta diversities are potentially useful in differentiating the types of contamination or identifying microbial sources of different origins.

**Fig. 2.**
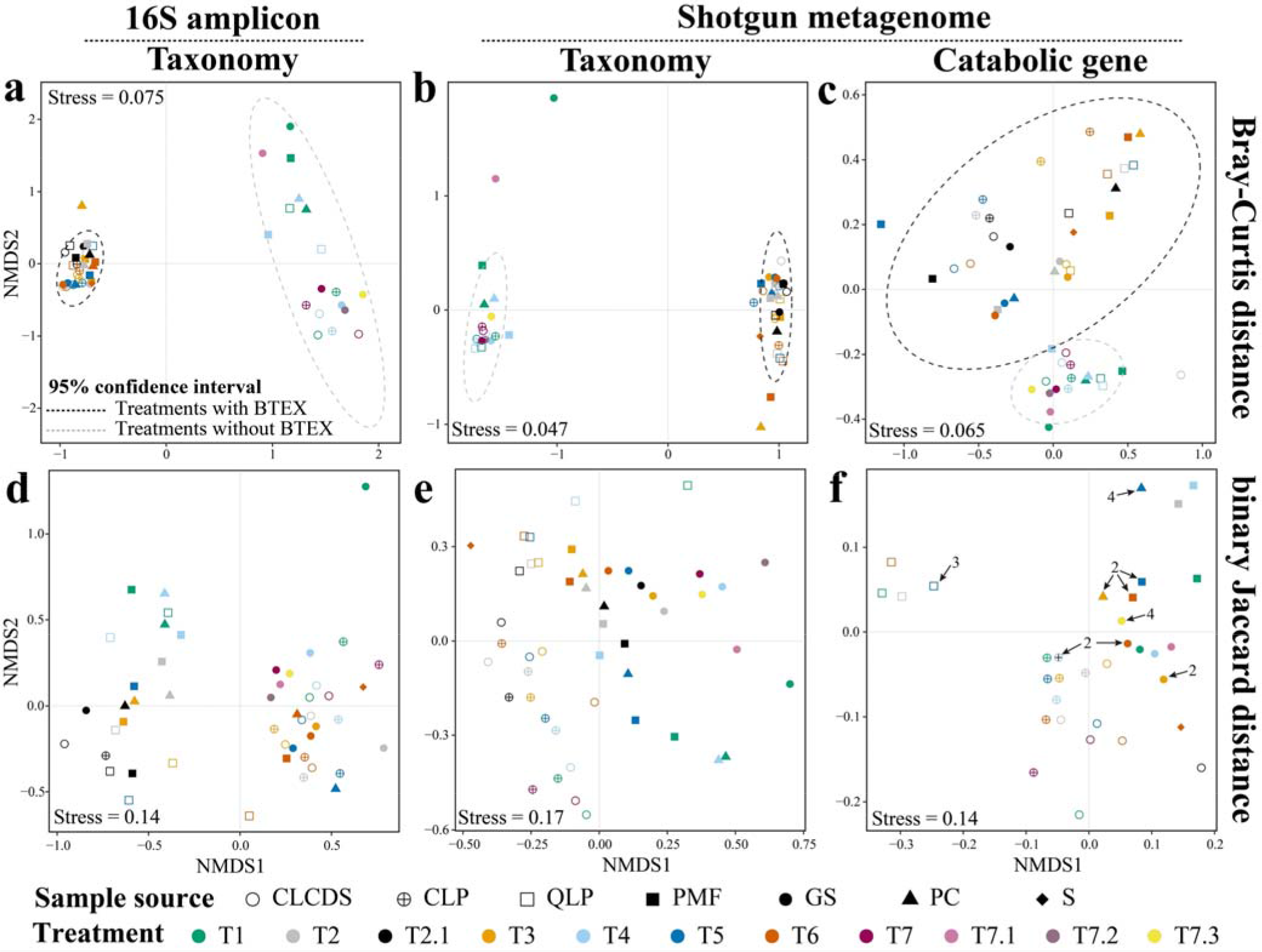
Microbial community composition differentiates among treatments, but metagenomic community gene diversity predicts inoculum sources, as indicated by non-metric multidimensional scaling (NMDS) ordination based on Bray-Curtis distance (plots a-c) and binary Jaccard distance (plots d-f) respectively (permutations = 9,999). (**a**) Genus profiles of 16S amplicon sequences, which was reanalyzed from the previously reported (Li et al. 2020) with three additional treatments (T7.1, T7.2, T7.3) from GS: Substrate (*p* < 1e-4, R^2^ = 0.817) + Source (*p* = 7e-03, R^2^ = 0.062); BTEX (presence versus absence) *p* < 1e-4, R^2^ = 0.729; Solvent (T2 versus T2.1) *p* = 0.11, R^2^ = 0.135. (**b**) Genus profiles of shotgun metagenome: Substrate (*p* < 1e-4, R^2^ = 0.816) + Source (*p* = 0.028, R^2^ = 0.055); BTEX (presence versus absence) *p* < 1e-4, R^2^ = 0.735; Solvent (T2 versus T2.1) *p* = 0.52, R^2^ = 0.064. (**c**) Community catabolic genes of shotgun metagenome: Substrate (*p* < 1e-4, R^2^ = 0.405) + Source (*p* = 0.0012, R^2^ = 0.198); BTEX (presence versus absence): *p* < 1e-4, R^2^ = 0.252; Solvent (T2 versus T2.1): *p* = 0.32, R^2^ = 0.095. (**d**) Genus profiles of 16S amplicon sequences: Substrate (*p* < 1e-4, R^2^ = 0.344) + Source (*p* < 1e-4, R^2^ = 0.213); BTEX (presence versus absence) *p* < 1e-4, R^2^ = 0.095; Solvent (T2 versus T2.1) *p* = 0.41, R^2^ = 0.077. (**e**) genus profiles of shotgun metagenome: Substrate (*p* < 1e-4, R^2^ = 0.244) + Source (*p* < 1e-4, R^2^ = 0.457); BTEX (presence versus absence) *p* < 1e-4, R^2^ = 0.113; Solvent (T2) versus T2.1 *p* = 0.89, R^2^ = 0.062. (**f**) community catabolic genes of shotgun metagenome: Substrate (*p* = 4e-04, R^2^ = 0.099) + Source (*p* < 1e-4, R^2^ = 0.790); BTEX (presence versus absence): *p* = 0.31, R^2^ = 0.025; Solvent (T2 versus T2.1): *p* = 0.79, R^2^ = 0.035. The R-squared and p-value are from multivariable permutational analysis of variance (PERMANOVA) using vegan adonis2 function in R. Shapes indicate sample sources of inocula; and colors indicate treatments with different xenobiotics. Dotted lines indicate the 95% confidence interval for the treatments in the presence (black line) and absence (gray line) of BTEX (plots a-c). Arrow indicates the points overlap; number indicates the number of the overlapped points (plot f). Chaka Lake Crystal Digging Site (CLCDS) sediment, Chaka Lake Port (CLP) sediment, Qinghai Lake Port (QLP) sediment, Gas Station (GS) subsurface soil, Petrochemical Complex (PC) soil, Plastic Manufacture Factory (PMF) soil, Solvent (S) contaminated site; T1: Dioxane (DMF), T2: BTEX (DMF), T2.1: BTEX (Dioxane), T3: BTEX + Dioxane (DMF), T4: Dioxane + TCE + TCA (DMF), T5: BTEX + TCE + TCA (DMF), T6: BTEX + Dioxane + TCE + TCA (DMF), T7: TCE + TCA (DMF), T7.1: TCA (DMF), T7.2: TCE (DMF), T7.3: PCE (DMF). See Table S1 for details of sample sources and treatments.

### 3.2. Diversity and phylogeny of xenobiotic degraders

To characterize the genes involved in the degradation of xenobiotics, such as BTEX (benzene, toluene, ethylbenzene, *o*-xylene, *m*-xylene, and *p*-xylene), perchloroethylene (PCE), trichloroethylene (TCE), trichloroethane (TCA), and vinyl chloride (VC), we analyzed the abundances of K numbers (KEGG Orthology identifiers) associated with KEGG metabolic pathways (map00361, map00623, map00642, map00622, and map00625) (Fig. S3), as well as *DMF* and K03418 for the small and large subunits of *N,N*-dimethylformamidase (DMFase) that break down the stable amide bond of the primary step of DMF degradation. 199 out of 1,052 genera harbor at least one of the target catabolic genes across all the enriched microbial communities (Fig. 3), accounting for the average of 94.6% of the microbial community compositions. Fungi were reported to possess diverse capacities to biodegrade organic contaminants (Harms et al. 2011), but only the genus *Ascochyta* harbors catabolic genes involved in the degradation of toluene and/or xylenes in samples (Fig. 3), suggesting bacteria are the major degraders of these xenobiotics. Bacterial xenobiotic degraders were primarily members of β- and γ-Proteobacteria and the Rhizobiales order within α-proteobacteria and one Actinobacteria genus *Rhodococcus* based on the catabolic gene abundance (Fig. 3). Some individual specific pathways appear to be confined to certain taxa. For example, across all enriched microbial communities, To2|B1 pathway genes were found in 9 genera; To3|B2 pathway genes were only found in *Paraburkholderia* and *Pseudomonas* genera as discussed below; K03380 for benzaldehyde dehydrogenase (To4 pathway) and K20199 for 4-hydroxybenzaldehyde dehydrogenase (To7 pathway) were only found in *Comamonas* and *Rhodococcus* genera. In addition, in the case of DMF metabolism, K03418 (i.e., the large subunit of DMFase) was identified in 6 genera (i.e., *Paracoccus*, *Hyphomicrobium*, *Achromobacter*, *Pontibaca*, *Starkeya*, and *Nitratireductor*), and the small subunit of DMFase in *Hyphomicrobium*, *Methyloversatilis*, and *Rhodopseudomonas* genera (Fig. 3, Fig. S4), which is further supported by the analyses of the *Paracoccus*, *Hyphomicrobium*, and *Achromobacter* MAGs (Fig. 4). The small subunit is likely to play a role in structural stabilization of the large subunit (Arya et al. 2020), but it is unclear why the small subunit is only present in certain genera. Averaged from all the enriched microbial communities in the treatments with BTEX, dominant genera (i.e., *Pseudomonas*, *Rhodococcus*, *Achromobacter*, *Alicycliphilus*, and *Comamonas*) generally carry relative highly abundant genes involved in the degradation of BTEX (Fig. 3). Low abundant taxa (e.g., *Thermomonas* and *Delftia*) may also carry relatively high abundant genes encoding for catabolic enzymes involved in some degradation pathways. Moreover, low abundant taxa may also contribute significantly to the xenobiotic degradation. For example, *Paraburkholderia* carries genes encoding enzymes for most of the degradation pathways, and particularly, together with *Pseudomonas*, are the only two genera harboring all genes associated with To3|B2 pathway (Fig. 3).

**Fig. 3.**
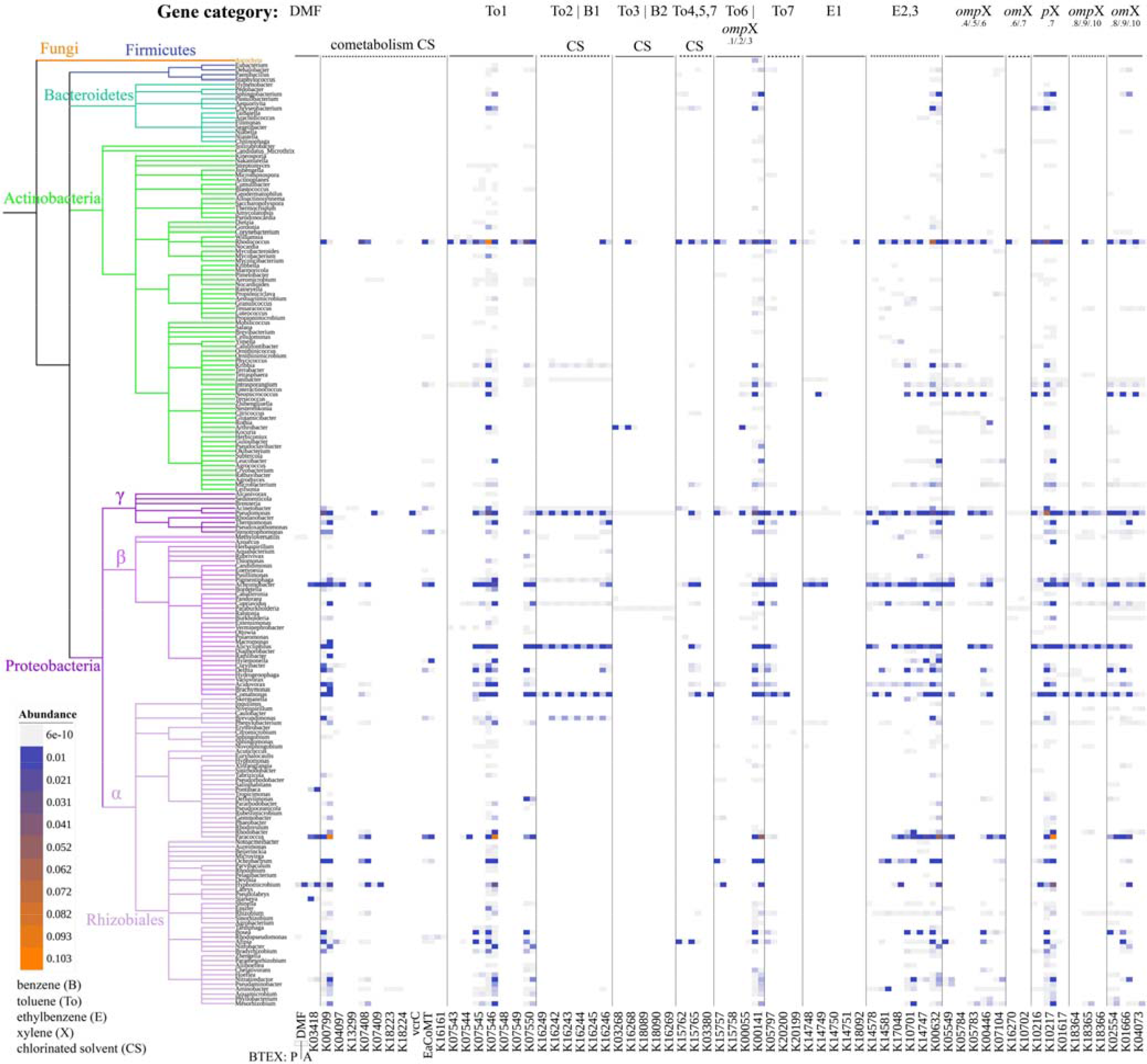
Phylogenetic distribution of catabolic genes involved in the xenobiotic degradation at the genus level. Each gene is shown in two columns to differentiate the treatments in the presence and absence of BTEX: in the left bottom, P indicates treatments in the presence of BTEX (in the left column of each gene); A indicates treatments in the absence of BTEX (in the right column of each gene). Genes are horizontally separated by specific pathways of xenobiotics. Color gradient indicates the gene abundance. This phylogenetic tree was obtained from NCBI common tree. See Fig. S4 for details of gene abundance of major genera in individual communities.

**Fig. 4.**
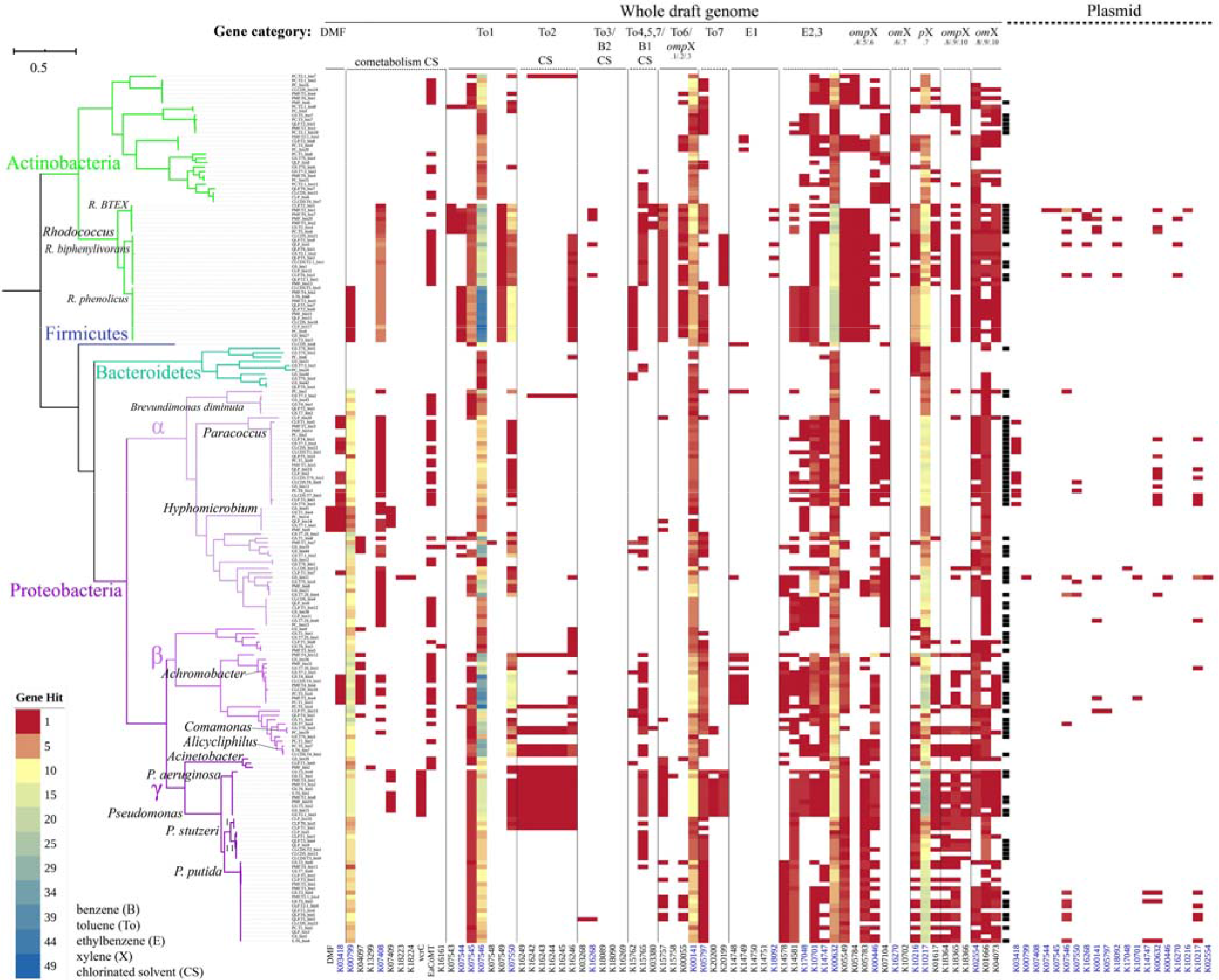
Metabolic potential of 201 metagenome-assembled genomes (> 70% complete, <5% contamination, < 99.8% similarity). The color gradient denotes the copy of genes associated with xenobiotic degradation in each genome. The column in black in the middle indicates plasmid-containing MAGs. The gene names in blue indicate at least one of the MAGs carries a plasmid harboring the gene. See Fig. S5 for taxonomy.

Across communities from different inoculum sources, the variation in catabolic pathway may occur among very closely related bacterial strains. For example, for the To2|B1 pathway, all catabolic genes were found in *Pseudomonas* enriched from CLP, GS, and PMF, partially CLCDS, but not from QLP and PC (Fig. S4). Additionally, genes involved in To3|B2 pathway and *om*X pathway (steps 6 and 7) were found in *Pseudomonas* enriched from PC and PMF but not from other inoculum sources, and for E1 pathway, only in GS *Pseudomonas* (Fig. S4).

### 3.3. Distribution patterns of catabolic genes in metagenomic assembled genomes

We recovered genomes by binning assembled and co-assembled contigs from 61 metagenomes, resulting in 201 metagenomic assembled genomes (MAGs) (on average, 93.3% completeness, 1.2% contamination). These MAGs represent four bacterial phyla, i.e., α-, β-, and γ-Proteobacteria, Actinobacteria, Bacteroidetes, and Firmicutes (Fig. S5), including 40 *Pseudomonas*, 32 *Rhodococcus*, 20 *Paracoccus*, 10 *Achromobacter*, 6 *Hyphomicrobium*, 3 *Alicycliphilus*, 3 *Acinetobacter*, and 2 *Comamonas* genomes (Fig. S5).

The catabolic genes vary not only among genera but also among species within the same genus. *Pseudomonas* species, i.e., *P*. *aeruginosa*, *P. stutzeri*, and *P. putida*, indicate distinct gene repertoires based on Bray-Curtis dissimilarity that accounts for gene presence and number (R^2^ = 0.725, *p* < 1e-4, PERMANOVA, Fig. S6a) and binary Jaccard dissimilarity that accounts for gene presence only (R^2^ = 0.633, *p* < 1e-4, PERMANOVA, Fig. S6c). For example, all *Pseudomonas* MAGs harbor K10217 (*p*X.7 pathway) but vary significantly among 3 species, with the highest copy number in *P*. *aeruginosa* (average 25.4), followed by *P. putida* (16.9), and *P. stutzeri* (8.2) (*p* < 0.0001, Kruskal-Wallis test, Fig. 4). In addition, K20200 and K20199 (To7 pathway) are present in *P*. *aeruginosa* but not in *P. stutzeri* and *P. putida* (Fig. 4). Likewise, *Rhodococcus* species (i.e., *R. BTEX*, *R. biphenylivorans*, and *R. phenolicus*) also reveal distinct gene repertoires (R^2^ = 0.862, *p* < 1e-4, Bray-Curtis distance, PERMANOVA, Fig. S6b; R^2^ = 0.809, *p* < 1e-4, binary Jaccard distance, PERMANOVA, Fig. S6d). For instance, K07543 (To1 pathway) was present in *R. BTEX* rather than *R. biphenylivorans* and *R. phenolicus*, whereas K16246 (To2 pathway) was present in *R. biphenylivorans* and *R. phenolicus* but *R. BTEX* (Fig. 4). These results suggest the microbial cooperation among different strains could play important roles in the complete degradation of aromatics.

### 3.4. Gene acquisition via plasmids appears to be common

To test our hypothesis of horizontal gene transfer (HGT) via plasmids, we predicted plasmids from MAGs and found 48.8% (98 out of 201) of the MAGs carry plasmid but differ considerably across genera. Among which, 100% of the *Paracoccus* MAGs were predicted to carry plasmid, and 45% for *Pseudomonas*, which is higher compared to the respective complete genomes deposited in NCBI (accessed May 11, 2020), where 90% for *Paracoccus* and 17.7% for *Pseudomonas* (Fig. 4, Table S2). This finding is consistent with the previous research demonstrating oil field discharges led to more plasmid-containing *Vibrio* strains and a greater number of plasmids per plasmid-containing strain (Hada and Sizemore 1981). The over one-year incubation in the presence of xenobiotics may result in the higher plasmid carriage rate of these two genera, whereas the species difference may also account for the observed difference. For example, none of the *Rhodococcus phenolicus* MAGs carry plasmids but all *Rhodococcus BTEX* MAGs carry plasmids, leading to a carriage rate of 37.5% for *Rhodococcus* MAGs, which is less than those complete genomes deposited in NCBI (66.7%). Neither the 6 *Hyphomicrobium* MAGs nor the 4 complete *Hyphomicrobium* genomes deposited in NCBI carry plasmids (Fig. 4, Table S2).

Twenty out of 67 genes involved in xenobiotic degradation were detected on plasmids across 201 MAGs (Fig. 4), which is consistent with the previous study showing that genes encoding enzymes involved in the degradation of organic contaminants are often located on plasmids (Garbisu et al. 2017). For instance, 13 out of 20 *Paracoccus* MAGs harbor K03418 encoding the large subunit of DMFase, among which 11 harbor K03418 on plasmids. Notably, 4 *Paracoccus* MAGs harbor one copy of K03418 on plasmids but chromosomes, whereas 7 harbor one copy on both chromosomes and plasmids (Fig. 4), suggesting HGT events in their acquisition of K03418 via plasmids in the microbial communities over one-year incubation. Likewise, 3 *P. putida* MAGs (i.e., GS.T3_bin4, GS.T5_bin5, and S.T6_bin4) harbor K14747 only on plasmids (Fig. 4). Multiple catabolic genes were also carried on the plasmids of *R. BTEX* and *R. biphenylivorans* strains. Nevertheless, gene acquisition via plasmids during long term enrichment could be limited to particular taxa, as not every strain carries a plasmid.

### 3.5. Efflux pumps enhance solvent tolerance

Efflux pumps can be the most effective resistance mechanism in response to the stress (Du et al. 2018). Like microbial antibiotic resistance, exposure to solvents can trigger the expression of transporters which provide efflux pathways, and therefore bacteria carrying efflux pumps can survive in the presence of toxic solvents (Du et al. 2018, Kusumawardhani et al. 2018, Udaondo et al. 2013). We estimated efflux pumps by searching against Resfams families (Gibson et al. 2015). Multiple families of efflux pumps were identified across the MAGs but were predominantly associated with the ATP-binding cassette (ABC), the major facilitator superfamily (MFS), and the resistance-nodulation-cell division (RND) families. A total of 29 efflux pump families were detected throughout the 201 MAGs, where high copies of RF0008 (ABC family) and RF0102 (MFS family) were ubiquitously observed (Fig. 5, Fig. S7). We validated these 29 efflux pump families via genomes of 4 major genera deposited in NCBI (Fig. S8). The copies of efflux pumps are within the reasonable range of individual genera, despite significant differences observed in some efflux pumps between MAGs of this study and NCBI genomes (10,352 *Pseudomonas*; 417 *Rhodococcus*; 132 *Paracoccus*; and 25 *Hyphomicrobium*).

**Fig. 5.**
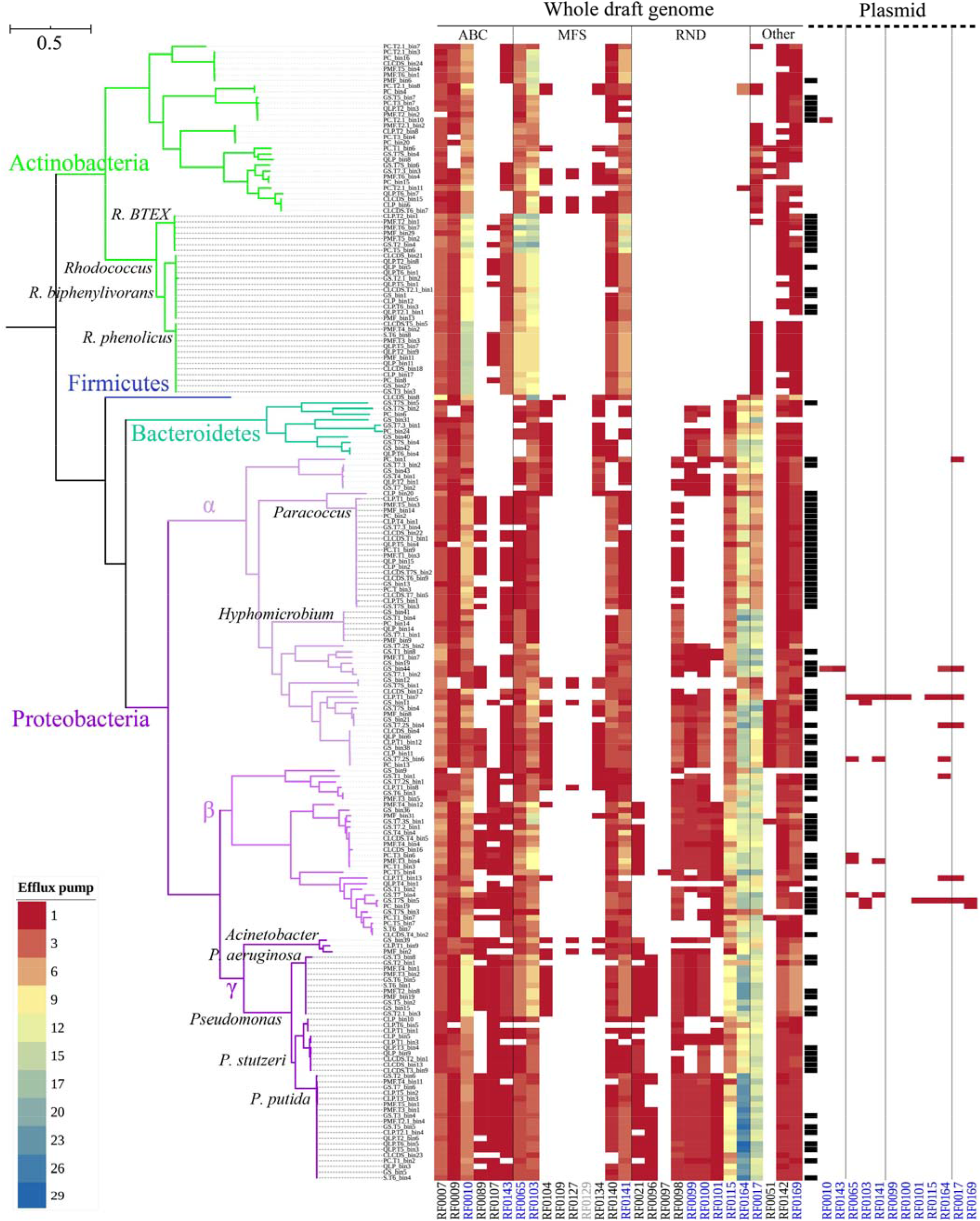
Efflux pumps of 201 MAGs. The color gradient denotes the copy of genes associated with efflux pumps in each genome. The column in black in the middle indicates plasmid-containing MAGs. The RF in blue in the bottom indicates efflux pumps detected in both plasmid and chromosome. RF0129 in grey was not detected in MAGs but NCBI genomes. See Fig. S7 for the same analysis including RF0008 and RF0102.

The efflux pumps of RND family were reported to be particularly important for the tolerance of *P. putida* to aromatics (Sol Cuenca et al. 2015), and *Pseudomonas* spp. have consistently higher copies of RND families except RF0098 among four major genera, irrespective of MAGs of this study or NCBI genomes (Fig. S8). Nine RND families were primarily found in MAGs of Proteobacteria and Bacteroidetes, which carry high copies of RF0164 (RND family). Additionally, the copies of RF0115 (RND family) in β- and γ-proteobacteria, particularly *P. putida* and *P*. *aeruginosa*, are high (Fig. 5). Comparatively, fewer gene copies of efflux pumps were observed in the MAGs of *P. stutzeri*, which were enriched from the sediments of saline lakes. *P*. *aeruginosa* MAGs also harbor relatively high copies of RF0010 (ABC family), while the highest is observed in *Rhodococcus*. Likewise, *Rhodococcus* harbor the highest copies of two MFS families (i.e., RF0065 and RF0103), albeit large variation occurs among *Rhodococcus* spp. Besides, *R. BTEX* MAGs harbor the highest copies of RF0141 (MFS family). In addition to the two ubiquitous efflux pumps, the copies of RF0164 and RF0017 efflux pumps are also high in *Hyphomicrobium*, and RF0164, high in *Paracoccus*. The efflux pumps could play a crucial role for bacterial survival in the presence of high concentrations of solvents.

Further, we evaluated the efflux pumps carried on plasmids. The carriage rates of the two ubiquitous efflux pumps on plasmids are high, i.e., 50% (49/98) for RF0008 and 37.7% (37/98) for RF0102 among the MAGs with plasmids, respectively (Fig. S7). Notably, all 20 *Paracoccus* MAGs harbor RF0008 on their carried plasmids, and 60% (12/20) *Paracoccus* for RF0102. Several other efflux pumps were also detected on plasmids of several MAGs, primarily in α- and β-proteobacteria. These results are suggestive of the acquisition of genes encoding efflux pumps via plasmids.

### 3.6. Co-occurrence network reveals predominant cooperation

We sought to dissect the microbial interactions within microbial communities. As the microbial communities are distinctive between treatments in the presence and absence of BTEX (Fig. 2), the microbial co-occurrence networks were constructed to represent treatments in the presence and absence of BTEX, using taxa that are present in at least half of the samples enriched with or without BTEX. Irrespective of treatments, the positive edges dominate interactions, i.e., 95% (presence) and 89% (absence) (Fig. 6, positive co-occurrence network; Fig. S9, negative co-occurrence network). In accordance with our hypothesis, xenobiotics are likely to promote synergistic interactions (Leewis et al. 2016). Thereafter, we focus on the positive networks, exhibiting high degrees of modularity (presence 0.863; absence 0.689), where 17 out of 48 modules account for 86.8% of the whole network nodes for treatments with BTEX (Fig. 6a), and for treatments without BTEX, 12 out of 32 modules account for 92.7% (Fig. 6b).

**Fig. 6.**
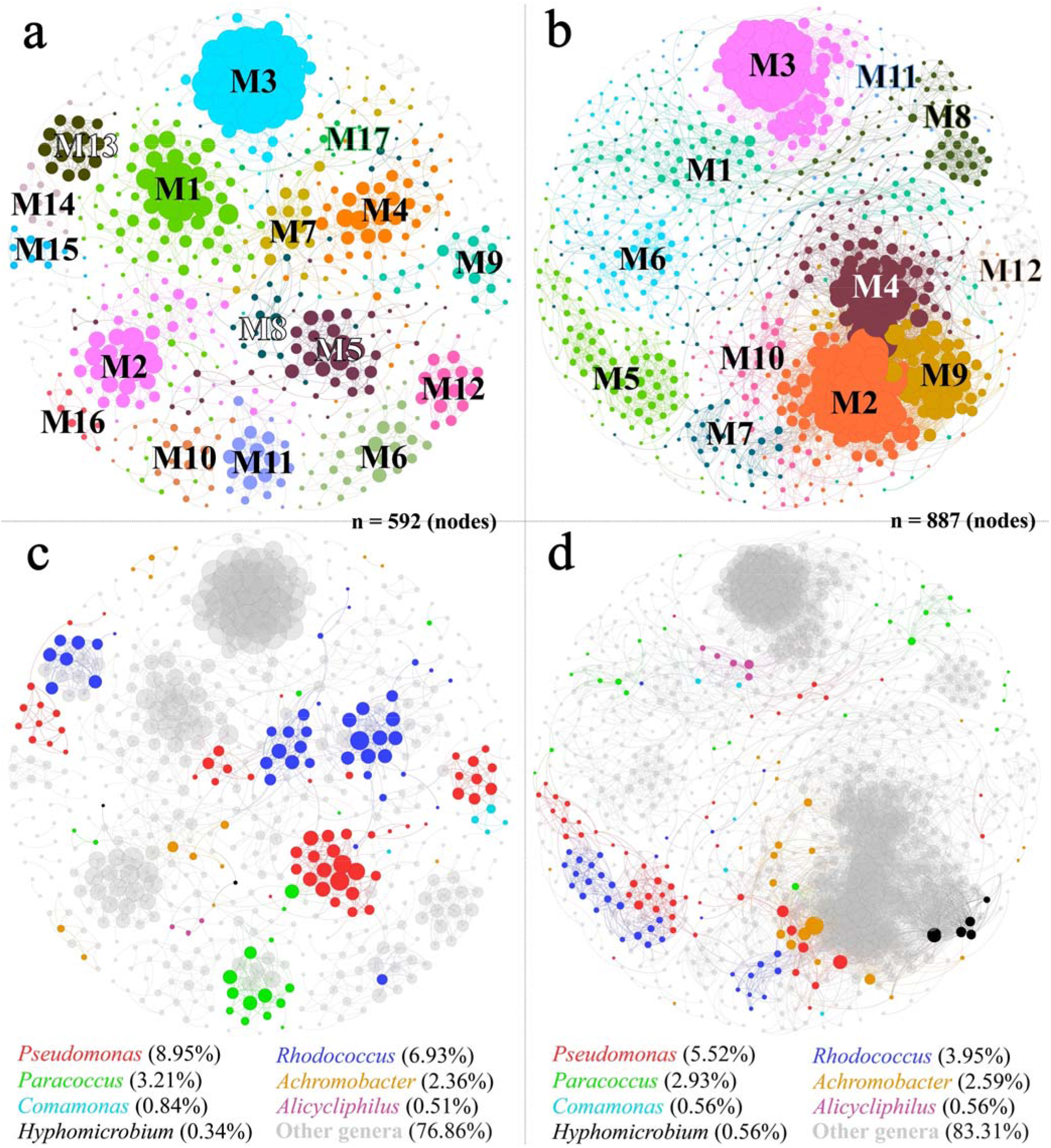
Positive microbial co-occurrence network among metagenomic taxa. **a**) Layout of major modules (> 1% of nodes, see Table S3a for details) and **c**) genus profiles for the treatments in the presence of BTEX; **b**) Layout of major modules (> 1% of nodes, see Table S3b for details) and **d**) genus profiles for the treatments in the absence of BTEX. A connection stands for a strong (Spearman ρ ◻>◻ 0.8) and significant (*p* < 0.01, multiple testing adjustment using Benjamini-Hochberg correction) correlation. Colors of plots **a** and **b**indicate modules; Colors of plots **c** and **d** indicate genera; Value in the bottom of plots **c** and **d**indicate node percentage of individual genera. Numbers on the figure indicate different modules. Circle size is proportional to the number of connections (i.e., degree) within each plot. See Fig. S9 for the negative co-occurrence network.

Distinct modules within microbial co-occurrence networks may reflect ecological niches (Lima-Mendez et al. 2015, Ma et al. 2020). The metagenomic microbial community composition differs significantly across modules based on genus-level taxonomy (R^2^ = 0.414 (presence), R^2^ = 0.422 (absence), *p* < 1e-4, PERMANOVA, Bray-Curtis distance; Fig. S10a,b; Table S4a). Likewise, the enriched microbial communities from different modules vary greatly in the catabolic gene composition (R^2^ = 0.339 (presence), R^2^ = 0.254 (absence), *p* < 1e-4, PERMANOVA, Bray-Curtis distance; Fig. S10c,d; Table S4b).

Major genera dominate the microbial communities but account for low proportions of nodes in the networks. Seven major genera account for 89.9% of the relative abundance across all metagenomes but account for 23.1% (presence, Fig. 6c) and 16.7% (absence, Fig. 6d) of nodes in the whole network respectively, and particularly *Paracoccus* and *Hyphomicrobium*, dominating treatments without BTEX (83.1%), account for 3.5% of nodes in the whole network (Fig. 6d). Nevertheless, the major genera are associated with multiple network modules, positively interacting with other minor genera. For example, in regards to the treatments with BTEX (Fig. 6c), *Pseudomonas*, accounting for 9.0% of the whole network nodes, are associated with five modules (1, 4, 5, 9, and 14). In Module 5, *Pseudomonas* strains dominate the network nodes (Fig. 6c), accounting for an average 36.3% of whole microbial community compositions of treatments in the presence of BTEX (Table S5), positively interacting with *Paracoccus*, *Comamonas* and *Alicycliphilus*, as well as strains of other minor genera. These results suggest that minor genera may play an important role in ecosystem functioning, albeit low abundant.

## 4. Discussion

Manipulating microbial communities is a key challenge facing bioremediation applications, and therefore is the primary motivation for this work. Recent studies have shown that environment turbulence (Hazen et al. 2010), introduction of microplastics (Seeley et al. 2020), ocean pollution (Chen et al. 2019), and resource pulses (i.e., glucose and/or ammonia) (Li et al. 2019) affect microbial community and functional gene composition. Through a combination of shotgun metagenomic sequencing and 16S rRNA gene amplicon sequencing, we characterized the microbiomes of 49 enriched microbial communities and recovered 201 MAGs involved in the degradation of xenobiotics. Insights from this work has enabled us to undercover the mechanisms underlying the success of major taxa in degrading xenobiotics.

The abundance of catabolic genes predicts the ability of microbes to degrade recalcitrant. In the microbial communities treated with BTEX, dominant taxa (e.g., *Pseudomonas*, *Rhodococcus*, *Achromobacter*, *Alicycliphilus*, and *Comamonas*) harbor catabolic genes that encode enzymes for multiple pathways of contaminant bioprocessing, whereas *Paracoccus* and *Hyphomicrobium* harbor genes encoding DMFase that dominate treatments without BTEX (Fig. 3, Fig. 4). Over 80% of the genera detected in the enriched microbial communities do not have any of the target genes, but they only account for a small proportion (5.4%) of the microbial community compositions. The intermediate metabolites produced by other taxa could be utilized as substrates for their growth.

The efflux pumps appear to play an essential role in aiding the survival of these microbes in the high concentrations of xenobiotics to overcome the toxicity of solvents and metabolites. All MAGs have multiple efflux pump families to tolerate xenobiotics. *Pseudomonas* spp. are highly tolerant to solvents, and several *P. putida* strains (i.e., DOT-T1E, S12, GM1, and MTB6) are able to thrive in the presence of high concentrations of aromatics, including BTEX (Rojo 2016, Sol Cuenca et al. 2015). *Pseudomonas* spp. harbor more genes encoding efflux pumps, e.g., *P. putida* and *P*. *aeruginosa* carry the highest copies of RF0164 and RF0115 across all MAGs. The efflux pumps are associated with the solvent tolerance but are not connected to the catabolic pathways (Sol Cuenca et al. 2015). Nevertheless, solvent tolerance traits are advantageous in microbial remediation of organic contaminants to overcome the toxicity of contaminants and products (Kusumawardhani et al. 2018), and thus solvent tolerance is an important property to consider in developing bioremediation systems (Sol Cuenca et al. 2015).

Plasmids which play an important role in bacterial evolution (Vial and Hommais 2020) can mediate HGT of catabolic genes (Nojiri et al. 2009, Shahi et al. 2017, Sun et al. 2017, Vedler 2009, Zhen et al. 2006). We predicted plasmids carried on the recovered MAGs and found 29.9% of the catabolic genes were harbored on plasmids in at least one of the MAGs. Intriguingly, certain genes (e.g., K03418 and K14747) were only located on the plasmid rather than the chromosome of some MAGs. These observations suggest HGT events in the acquisition of catabolic genes via plasmids in the microbial communities over a one-year incubation. Microorganisms are able to develop tolerance mechanism once exposed to solvents (van der Meer 2006), and plasmids have been implicated in the spread of solvent-extruding efflux pumps (Rodriguez-Herva et al. 2007, Segura et al. 2012, Sol Cuenca et al. 2015). Nearly half (14/29) of the observed efflux pump families in the MAGs were found on plasmids, and approximately 30% (60/201) of MAGs harbor at least one efflux pump family on their plasmids (Fig. S7). Plasmids contribute to genome plasticity (Dobrindt and Hacker 2001), facilitating rapid evolution and adaptation of their hosts for the degradation of xenobiotics by acquiring genes involved in metabolism and tolerance. However, plasmid carriage varies considerably among genera, e.g., of two major genera in the treatments without BTEX, all *Paracoccus* strains carry plasmids but none of the *Hyphomicrobium* strains carries plasmids. The plasmid-containing strains have some advantages (e.g., acquiring genes which complement chromosomally encoded functions in metabolism and solvent tolerance) over the non-plasmid-containing strains, which is another critical property to consider for the development of bioremediation systems.

Our findings also emphasize the synergistic interactions among the enriched microbial communities. It will be beneficial to apply microbial communities in the xenobiotic biodegradation. Complex recalcitrant substrates also promote positive cooperation among mixed cultures (Cortes-Tolalpa et al. 2017, Deng and Wang 2017, Leewis et al. 2016), as the complete mineralization of recalcitrant substrates requires the complementary interaction of a set of enzymes. The interaction among microbial communities depends on the recalcitrance of xenobiotics. For example, in the case of *m*-xylene, the biotransformation to propanoyl-CoA is associated with over 10 enzymes (Fig. S3). Therefore, the combinations of microbial taxa efficiently degrade *m*-xylene, releasing small molecular substrates. As anticipated, major genera are associated with multiple co-occurrence network modules, among which, most modules consist of different taxa from multiple genera, including both major and minor genera. The taxa with lower abundance may fulfill essential functions in the degradation and enhance functionality of the abundant microbes (Jousset et al. 2017, Xue et al. 2018).

## 5. Conclusions

The integration of multiple microbial traits of microbial community assembly has rarely been considered in microbial degradation of contaminants, although individual microbial traits, particularly catabolic traits, have been frequently studied for microbial degradation. Results presented here show how the integration of catabolic traits, solvent tolerance traits, plasmid-mediated gene acquisition, and synergistic interactions among the enriched microbial communities advance our understanding of microbial processes driving the biodegradation of xenobiotics. Our study provides a framework for the incorporation of trait-based microbial strategies into the xenobiotic degradation. The findings may have important implications for the development of bioremediation systems.

## Acknowledgements

This work was supported by the National Natural Science Foundation of China (Grant No. 51409106) and the University of Macau Multi-Year Research Grant (MYRG2018-00108-FST). We thank Karissa Lynn Cross for feedback on previous versions of the manuscript and Xia Huang, Shuhuan Wang, Jinhuan Liu, Yingshi Li, Jiying Liu, and Haimei Su for their assistance in the laboratory.

## References

Alneberg, J., Bjarnason, B.S., de Bruijn, I., Schirmer, M., Quick, J., Ijaz, U.Z., Lahti, L., Loman, N.J., Andersson, A.F. and Quince, C., 2014. Binning metagenomic contigs by coverage and composition. Nature Methods 11(11), 1144–1146.

Aramaki, T., Blanc-Mathieu, R., Endo, H., Ohkubo, K., Kanehisa, M., Goto, S. and Ogata, H., 2020. KofamKOALA: KEGG Ortholog assignment based on profile HMM and adaptive score threshold. Bioinformatics 36(7), 2251–2252.

Arya, C.K., Yadav, S., Fine, J., Casanal, A., Chopra, G., Ramanathan, G., Vinothkumar, K.R. and Subramanian, R., 2020. Breaking a Strong Amide Bond: Structure and Properties of Dimethylformamidase. bioRxiv, 2019.2012.2017.879908.

Bastian, M., Heymann, S. and Jacomy, M., 2009. Gephi: an open source software for exploring and manipulating networks.

Basu, A., Apte, S.K. and Phale, P.S., 2006. Preferential utilization of aromatic compounds over glucose by Pseudomonas putida CSV86. Applied and environmental microbiology 72(3), 2226.

Ben Said, S. and Or, D., 2017. Synthetic microbial ecology: Engineering habitats for modular consortia. Frontiers in Microbiology 8, 1125–1125.

Berlemont, R. and Martiny, A.C., 2015. Genomic potential for polysaccharide deconstruction in bacteria. Applied and environmental microbiology 81(4), 1513.

Blondel, V., Guillaume, J.-L., Lambiotte, R. and Lefebvre, E., 2008. Fast Unfolding of Communities in Large Networks. Journal of Statistical Mechanics Theory and Experiment 2008.

Capella-Gutiérrez, S., Silla-Martínez, J.M. and Gabaldón, T., 2009. trimAl: a tool for automated alignment trimming in large-scale phylogenetic analyses. Bioinformatics (Oxford, England) 25(15), 1972–1973.

Cerqueira, V.S., Hollenbach, E.B., Maboni, F., Vainstein, M.H., Camargo, F.A.O., Peralba, M.d.C.R. and Bento, F.M., 2011. Biodegradation potential of oily sludge by pure and mixed bacterial cultures. Bioresource Technology 102(23), 11003–11010.

Chen, J., McIlroy, S.E., Archana, A., Baker, D.M. and Panagiotou, G., 2019. A pollution gradient contributes to the taxonomic, functional, and resistome diversity of microbial communities in marine sediments. Microbiome 7(1), 104.

Cortes-Tolalpa, L., Salles, J.F. and van Elsas, J.D., 2017. Bacterial synergism in lignocellulose biomass degradation - complementary roles of degraders as influenced by complexity of the carbon source. Frontiers in Microbiology 8(1628).

de Lima-Morales, D., Chaves-Moreno, D., Wos-Oxley, M.L., Jáuregui, R., Vilchez-Vargas, R. and Pieper, D.H., 2016. Degradation of benzene by Pseudomonas veronii 1YdBTEX2 and 1YB2 is catalyzed by enzymes encoded in distinct catabolism gene clusters. Applied and environmental microbiology 82(1), 167–173.

Deng, Y.-J. and Wang, S.Y., 2017. Complex carbohydrates reduce the frequency of antagonistic interactions among bacteria degrading cellulose and xylan. FEMS Microbiology Letters 364(5), fnx019–fnx019.

Dobrindt, U. and Hacker, J., 2001. Whole genome plasticity in pathogenic bacteria. Current Opinion in Microbiology 4(5), 550–557.

Du, D., Wang-Kan, X., Neuberger, A., van Veen, H.W., Pos, K.M., Piddock, L.J.V. and Luisi, B.F., 2018. Multidrug efflux pumps: structure, function and regulation. Nature Reviews Microbiology 16(9), 523–539.

Faust, K., 2019. Microbial consortium design benefits from metabolic modeling. Trends in Biotechnology 37(2), 123–125.

Finn, R.D., Clements, J. and Eddy, S.R., 2011. HMMER web server: interactive sequence similarity searching. Nucleic Acids Research 39(suppl_2), W29–W37.

Garbisu, C., Garaiyurrebaso, O., Epelde, L., Grohmann, E. and Alkorta, I., 2017. Plasmid-mediated bioaugmentation for the bioremediation of contaminated soils. Frontiers in Microbiology 8, 1966–1966.

Garcia, S., 2016. Mixed cultures as model communities: Hunting for ubiquitous microorganisms, their partners, and interactions. Aquatic Microbial Ecology 77.

Gibson, M.K., Forsberg, K.J. and Dantas, G., 2015. Improved annotation of antibiotic resistance determinants reveals microbial resistomes cluster by ecology. The ISME Journal 9(1), 207–216.

Hada, H.S. and Sizemore, R.K., 1981. Incidence of plasmids in marine Vibrio spp. isolated from an oil field in the northwestern Gulf of Mexico. Applied and environmental microbiology 41(1), 199–202.

Hall, J.P.J., Wood, A.J., Harrison, E. and Brockhurst, M.A., 2016. Source-sink plasmid transfer dynamics maintain gene mobility in soil bacterial communities. Proceedings of the National Academy of Sciences of the United States of America 113(29), 8260–8265.

Harms, H., Schlosser, D. and Wick, L.Y., 2011. Untapped potential: exploiting fungi in bioremediation of hazardous chemicals. Nature Reviews Microbiology 9(3), 177–192.

Hays, S.G., Patrick, W.G., Ziesack, M., Oxman, N. and Silver, P.A., 2015. Better together: engineering and application of microbial symbioses. Current Opinion in Biotechnology 36, 40–49.

Hazen, T.C., Dubinsky, E.A., DeSantis, T.Z., Andersen, G.L., Piceno, Y.M., Singh, N., Jansson, J.K., Probst, A., Borglin, S.E., Fortney, J.L., Stringfellow, W.T., Bill, M., Conrad, M.E., Tom, L.M., Chavarria, K.L., Alusi, T.R., Lamendella, R., Joyner, D.C., Spier, C., Baelum, J., Auer, M., Zemla, M.L., Chakraborty, R., Sonnenthal, E.L., D’haeseleer, P., Holman, H.-Y.N., Osman, S., Lu, Z., Van Nostrand, J.D., Deng, Y., Zhou, J. and Mason, O.U., 2010. Deep-Sea Oil Plume Enriches Indigenous Oil-Degrading Bacteria. Science 330(6001), 204.

Hyatt, D., Chen, G.-L., Locascio, P.F., Land, M.L., Larimer, F.W. and Hauser, L.J., 2010. Prodigal: prokaryotic gene recognition and translation initiation site identification. BMC bioinformatics 11, 119–119.

Inoue, A. and Horikoshi, K., 1989. A Pseudomonas thrives in high concentrations of toluene. Nature 338(6212), 264–266.

Isobe, K., Allison, S.D., Khalili, B., Martiny, A.C. and Martiny, J.B.H., 2019. Phylogenetic conservation of bacterial responses to soil nitrogen addition across continents. Nature Communications 10(1), 2499.

Jousset, A., Bienhold, C., Chatzinotas, A., Gallien, L., Gobet, A., Kurm, V., Küsel, K., Rillig, M.C., Rivett, D.W., Salles, J.F., van der Heijden, M.G.A., Youssef, N.H., Zhang, X., Wei, Z. and Hol, W.H.G., 2017. Where less may be more: how the rare biosphere pulls ecosystems strings. The ISME Journal 11(4), 853–862.

Kang, D., Jacquiod, S., Herschend, J., Wei, S., Nesme, J. and Sørensen, S.J., 2020. Construction of simplified microbial consortia to degrade recalcitrant materials based on enrichment and dilution-to-extinction cultures. Frontiers in Microbiology 10(3010).

Kang, D.D., Li, F., Kirton, E., Thomas, A., Egan, R., An, H. and Wang, Z., 2019. MetaBAT 2: an adaptive binning algorithm for robust and efficient genome reconstruction from metagenome assemblies. PeerJ 7, e7359–e7359.

Krause, S., Le Roux, X., Niklaus, P.A., Van Bodegom, P.M., Lennon, J.T., Bertilsson, S., Grossart, H.-P., Philippot, L. and Bodelier, P.L.E., 2014. Trait-based approaches for understanding microbial biodiversity and ecosystem functioning. Frontiers in Microbiology 5, 251–251.

Krueger, F., 2015. Trim galore. A wrapper tool around Cutadapt and FastQC to consistently apply quality and adapter trimming to FastQ files 516, 517.

Kusumawardhani, H., Hosseini, R. and de Winde, J.H., 2018. Solvent tolerance in bacteria: fulfilling the promise of the biotech era? Trends in Biotechnology 36(10), 1025–1039.

Lajoie, G. and Kembel, S.W., 2019. Making the Most of Trait-Based Approaches for Microbial Ecology. Trends in Microbiology 27(10), 814–823.

Langmead, B. and Salzberg, S.L., 2012. Fast gapped-read alignment with Bowtie 2. Nature methods 9(4), 357–359.

Leewis, M.-C., Uhlik, O. and Leigh, M.B., 2016. Synergistic processing of biphenyl and benzoate: carbon flow through the bacterial community in polychlorinated-biphenyl-contaminated soil. Scientific Reports 6, 22145–22145.

Letunic, I. and Bork, P., 2016. Interactive tree of life (iTOL) v3: an online tool for the display and annotation of phylogenetic and other trees. Nucleic Acids Research 44(W1), W242–W245.

Li, D., Liu, C.-M., Luo, R., Sadakane, K. and Lam, T.-W., 2015. MEGAHIT: an ultra-fast single-node solution for large and complex metagenomics assembly via succinct de Bruijn graph. Bioinformatics 31(10), 1674–1676.

Li, H. and Durbin, R., 2009. Fast and accurate short read alignment with Burrows-Wheeler transform. Bioinformatics (Oxford, England) 25(14), 1754–1760.

Li, H., Handsaker, B., Wysoker, A., Fennell, T., Ruan, J., Homer, N., Marth, G., Abecasis, G., Durbin, R. and Genome Project Data Processing, S., 2009. The Sequence Alignment/Map format and SAMtools. Bioinformatics 25(16), 2078–2079.

Li, J., Mau, R.L., Dijkstra, P., Koch, B.J., Schwartz, E., Liu, X.-J.A., Morrissey, E.M., Blazewicz, S.J., Pett-Ridge, J., Stone, B.W., Hayer, M. and Hungate, B.A., 2019. Predictive genomic traits for bacterial growth in culture versus actual growth in soil. The ISME Journal 13, 2162–2172.

Li, J., Wu, C., Chen, S., Lu, Q., Shim, H., Huang, X., Jia, C. and Wang, S., 2020. Enriching indigenous microbial consortia as a promising strategy for xenobiotics’ cleanup. Journal of Cleaner Production 261, 121234.

Lima-Mendez, G., Faust, K., Henry, N., Decelle, J., Colin, S., Carcillo, F., Chaffron, S., Ignacio-Espinosa, J.C., Roux, S., Vincent, F., Bittner, L., Darzi, Y., Wang, J., Audic, S., Berline, L., Bontempi, G., Cabello, A.M., Coppola, L., Cornejo-Castillo, F.M., Ovidio, F., De Meester, L., Ferrera, I., Garet-Delmas, M.-J., Guidi, L., Lara, E., Pesant, S., Royo-Llonch, M., Salazar, G., Sánchez, P., Sebastian, M., Souffreau, C., Dimier, C., Picheral, M., Searson, S., Kandels-Lewis, S., Gorsky, G., Not, F., Ogata, H., Speich, S., Stemmann, L., Weissenbach, J., Wincker, P., Acinas, S.G., Sunagawa, S., Bork, P., Sullivan, M.B., Karsenti, E., Bowler, C., de Vargas, C. and Raes, J., 2015. Determinants of community structure in the global plankton interactome. Science 348(6237), 1262073.

Lindemann, S.R., Bernstein, H.C., Song, H.-S., Fredrickson, J.K., Fields, M.W., Shou, W., Johnson, D.R. and Beliaev, A.S., 2016. Engineering microbial consortia for controllable outputs. The ISME Journal 10(9), 2077–2084.

Ma, B., Wang, Y., Ye, S., Liu, S., Stirling, E., Gilbert, J.A., Faust, K., Knight, R., Jansson, J.K., Cardona, C., Röttjers, L. and Xu, J., 2020. Earth microbial co-occurrence network reveals interconnection pattern across microbiomes. Microbiome 8(1), 82.

Malik, A.A., Martiny, J.B.H., Brodie, E.L., Martiny, A.C., Treseder, K.K. and Allison, S.D., 2020. Defining trait-based microbial strategies with consequences for soil carbon cycling under climate change. The ISME Journal 14(1), 1–9.

Martiny, J.B.H., Jones, S.E., Lennon, J.T. and Martiny, A.C., 2015. Microbiomes in light of traits: A phylogenetic perspective. Science 350(6261), aac9323.

Mende, D.R., Sunagawa, S., Zeller, G. and Bork, P., 2013. Accurate and universal delineation of prokaryotic species. Nature Methods 10(9), 881–884.

Menzel, P., Ng, K.L. and Krogh, A., 2016. Fast and sensitive taxonomic classification for metagenomics with Kaiju. Nature Communications 7(1), 11257.

Minh, B.Q., Schmidt, H.A., Chernomor, O., Schrempf, D., Woodhams, M.D., von Haeseler, A. and Lanfear, R., 2020. IQ-TREE 2: new models and efficient methods for phylogenetic inference in the genomic Era. Molecular Biology and Evolution 37(5), 1530–1534.

Molina-Santiago, C., Udaondo, Z., Gómez-Lozano, M., Molin, S. and Ramos, J.-L., 2017. Global transcriptional response of solvent-sensitive and solvent-tolerant Pseudomonas putida strains exposed to toluene. Environmental Microbiology 19(2), 645–658.

Nojiri, H., Sota, M. and Shintani, M., 2009. Microbial Megaplasmids. Schwartz, E. (ed), pp. 55–87, Springer Berlin Heidelberg, Berlin, Heidelberg.

Oksanen, J., Blanchet, F.G., Friendly, M., Kindt, R., Legendre, P., McGlinn, D., Minchin, P.R., O’Hara, R.B., Simpson, G.L., Solymos, P., Henry, M., Stevens, H., Szoecs, E. and Wagner, H., 2013. Package ‘vegan’. Community ecology package 2(9).

Olm, M.R., Brown, C.T., Brooks, B. and Banfield, J.F., 2017. dRep: a tool for fast and accurate genomic comparisons that enables improved genome recovery from metagenomes through de-replication. The ISME Journal 11(12), 2864–2868.

Parks, D.H., Imelfort, M., Skennerton, C.T., Hugenholtz, P. and Tyson, G.W., 2015. CheckM: assessing the quality of microbial genomes recovered from isolates, single cells, and metagenomes. Genome Res 25(7), 1043–1055.

Poursat, B.A.J., van Spanning, R.J.M., de Voogt, P. and Parsons, J.R., 2019. Implications of microbial adaptation for the assessment of environmental persistence of chemicals. Critical Reviews in Environmental Science and Technology 49(23), 2220–2255.

Rodriguez-Herva, J., Garcia, V., Hurtado, A., Segura, A. and Ramos, J., 2007. The ttgGHI solvent efflux pump operon of Pseudomonas putida DOT-T1E Is located on a large self-transmissible plasmid. Environmental Microbiology 9, 1550–1561.

Rojo, F., 2016. Traits allowing resistance to organic solvents in Pseudomonas. Environmental Microbiology 19(2), 417–419.

Schwengers, O., Barth, P., Falgenhauer, L., Hain, T., Chakraborty, T. and Goesmann, A., 2020. Platon: identification and characterization of bacterial plasmid contigs in short-read draft assemblies exploiting protein-sequence-based replicon distribution scores. bioRxiv, 2020.2004.2021.053082.

Seeley, M.E., Song, B., Passie, R. and Hale, R.C., 2020. Microplastics affect sedimentary microbial communities and nitrogen cycling. Nature Communications 11(1), 2372.

Segura, A., Molina, L., Fillet, S., Krell, T., Bernal, P., Muñoz-Rojas, J. and Ramos, J.-L., 2012. Solvent tolerance in Gram-negative bacteria. Current Opinion in Biotechnology 23(3), 415–421.

Shahi, A., Ince, B., Aydin, S. and Ince, O., 2017. Assessment of the horizontal transfer of functional genes as a suitable approach for evaluation of the bioremediation potential of petroleum-contaminated sites: a mini-review. Applied Microbiology and Biotechnology 101(11), 4341–4348.

Sievers, F. and Higgins, D.G., 2014. Clustal Omega. Current Protocols in Bioinformatics 48(1), 3.13.11–13.13.16.

Sievers, F., Wilm, A., Dineen, D., Gibson, T.J., Karplus, K., Li, W., Lopez, R., McWilliam, H., Remmert, M., Söding, J., Thompson, J.D. and Higgins, D.G., 2011. Fast, scalable generation of high-quality protein multiple sequence alignments using Clustal Omega. Mol Syst Biol 7, 539–539.

Singh, A. and Ward, O.P., 2004. Biodegradation and Bioremediation. Singh, A. and Ward, O.P. (eds), pp. 1–17, Springer Berlin Heidelberg, Berlin, Heidelberg.

Sol Cuenca, M., Gómez-García, M.R., Udaondo, Z., Segura, A., Molina-Santiago, C., Duque, E., Ramos, J.-L. and Roca, A., 2015. Mechanisms of solvent resistance mediated by interplay of cellular factors in Pseudomonas putida. FEMS Microbiology Reviews 39(4), 555–566.

Sorensen, J.W., Dunivin, T.K., Tobin, T.C. and Shade, A., 2019. Ecological selection for small microbial genomes along a temperate-to-thermal soil gradient. Nature Microbiology 4(1), 55–61.

Sun, J., Qiu, Y., Ding, P., Peng, P., Yang, H. and Li, L., 2017. Conjugative transfer of dioxin-catabolic megaplasmids and bioaugmentation prospects of a Rhodococcus sp. Environmental Science & Technology 51(11), 6298–6307.

Suttinun, O., Luepromchai, E. and Müller, R., 2013. Cometabolism of trichloroethylene: concepts, limitations and available strategies for sustained biodegradation. Reviews in Environmental Science and Bio/Technology 12(1), 99–114.

Udaondo, Z., Molina, L., Daniels, C., Gómez, M.J., Molina-Henares, M.A., Matilla, M.A., Roca, A., Fernández, M., Duque, E., Segura, A. and Ramos, J.L., 2013. Metabolic potential of the organic-solvent tolerant Pseudomonas putida DOT-T1E deduced from its annotated genome. Microbial Biotechnology 6(5), 598–611.

Uritskiy, G.V., DiRuggiero, J. and Taylor, J., 2018. MetaWRAP—a flexible pipeline for genome-resolved metagenomic data analysis. Microbiome 6(1), 158.

van der Meer, J.R., 2006. Environmental pollution promotes selection of microbial degradation pathways. Frontiers in Ecology and the Environment 4(1), 35–42.

van der Meer, J.R., de Vos, W.M., Harayama, S. and Zehnder, A.J., 1992. Molecular mechanisms of genetic adaptation to xenobiotic compounds. Microbiol Rev 56(4), 677–694.

Vedler, E., 2009. Microbial Megaplasmids. Schwartz, E. (ed), pp. 33–53, Springer Berlin Heidelberg, Berlin, Heidelberg.

Vial, L. and Hommais, F., 2020. Plasmid-chromosome cross-talks. Environmental Microbiology 22(2), 540–556.

Wu, D., Jospin, G. and Eisen, J.A., 2013. Systematic identification of gene families for use as “markers” for phylogenetic and phylogeny-driven ecological studies of bacteria and archaea and their major subgroups. PLOS ONE 8(10), e77033.

Wu, Y.-W., Simmons, B.A. and Singer, S.W., 2016. MaxBin 2.0: an automated binning algorithm to recover genomes from multiple metagenomic datasets. Bioinformatics 32(4), 605–607.

Xu, X., Zarecki, R., Medina, S., Ofaim, S., Liu, X., Chen, C., Hu, S., Brom, D., Gat, D., Porob, S., Eizenberg, H., Ronen, Z., Jiang, J. and Freilich, S., 2019. Modeling microbial communities from atrazine contaminated soils promotes the development of biostimulation solutions. The ISME Journal 13(2), 494–508.

Xue, Y., Chen, H., Yang, J.R., Liu, M., Huang, B. and Yang, J., 2018. Distinct patterns and processes of abundant and rare eukaryotic plankton communities following a reservoir cyanobacterial bloom. The ISME Journal 12(9), 2263–2277.

Zhen, D., Liu, H., Wang, S.-J., Zhang, J.-J., Zhao, F. and Zhou, N.-Y., 2006. Plasmid-mediated degradation of 4-chloronitrobenzene by newly isolated Pseudomonas putida strain ZWL73. Applied Microbiology and Biotechnology 72(4), 797–803.

Zhou, X., Jin, W., Sun, C., Gao, S.-H., Chen, C., Wang, Q., Han, S.-F., Tu, R., Latif, M.A. and Wang, Q., 2018. Microbial degradation of N, N-dimethylformamide by Paracoccus sp. strain DMF-3 from activated sludge. Chemical Engineering Journal 343, 324–330.

