## Supplemental files for "Mechanistic insights into the success of xenobiotic degraders resolved from metagenomes of microbial enrichment cultures"

*Bioinformatic analyses of 16S rRNA gene amplicon sequencing*

Data processing and analyses of 16S rRNA gene amplicon sequencing were performed as described previously (Li et al. 2020). In short, the DNA quality and quantity were estimated using the Implen NanoDrop (Munich, Germany). The 16S rRNA genes (V4-V5 region) were amplified and sequenced using Illumina Miseq sequencing (Illumina, San Diego, CA, USA). The provided pair-end (250 × 2) demultiplexed sequences were assembled and filtered using DADA2 (Callahan et al. 2016). The resulting amplicon sequencing variants (ASVs) were compared against the SILVA taxonomic training data formatted for DADA2 (SILVA v132). Samples were rarefied to 10,000 reads 100 times, then the average counts of all 100 rarefied tables for the downstream analysis were calculated.

*Concordance between shotgun metagenome and 16S amplicon sequencing data*

Amplicon sequencing of a region of 16S rRNA gene often does not provide effective resolution below the genus level (Yarza et al. 2014), and hence we compared the overall concordance between shotgun metagenomic and 16S amplicon sequencing genus profiles for pairs of each enriched sample. Eighty-four genera are present in at least one of the enriched microbial communities for both shotgun metagenomic and 16S amplicon sequencing profiles, accounting for 93.5% (80.1% – 97.9%) of shotgun metagenomic sequences and 97.2% (92.0% – 98.9%) of 16S amplicon sequences, respectively. The Pearson correlation R-squared value was 0.91 (*p* < 2.2e-16, Fig. S1a) across all enriched sample pairs, with an average of 0.91 (*p* = 3.4e-10, Fig. S1b) of individual enriched microbial communities. Among the nine major genera (i.e., 7 metagenomic genera with the average relative abundance ≥ 1% in the treatments in the presence or absence of BTEX; 2 genera from amplicon sequencing analyses), in general, most are comparatively abundant between sequencing methods, except *Chryseobacterium* and *Sphingobacterium*, which are much less abundant in metagenome analyses as compared to 16S amplicon analyses (Fig. S2). The discrepancy could be attributed to multiple factors, e.g., library preparation methods (Gohl et al. 2016), regions of 16S gene (Johnson et al. 2019, Rausch et al. 2019), analytical pipelines (Prodan et al. 2020), and different copy numbers of 16S rRNA gene (Louca et al. 2018). Nevertheless, taken together with comparable variation explained by substrate and inoculum source in microbial community compositions from the analyses as demonstrated (Fig. 2a,b), we conclude that in general amplicon and metagenomic sequencing approaches reveal consistent taxa in the enriched microbial communities.

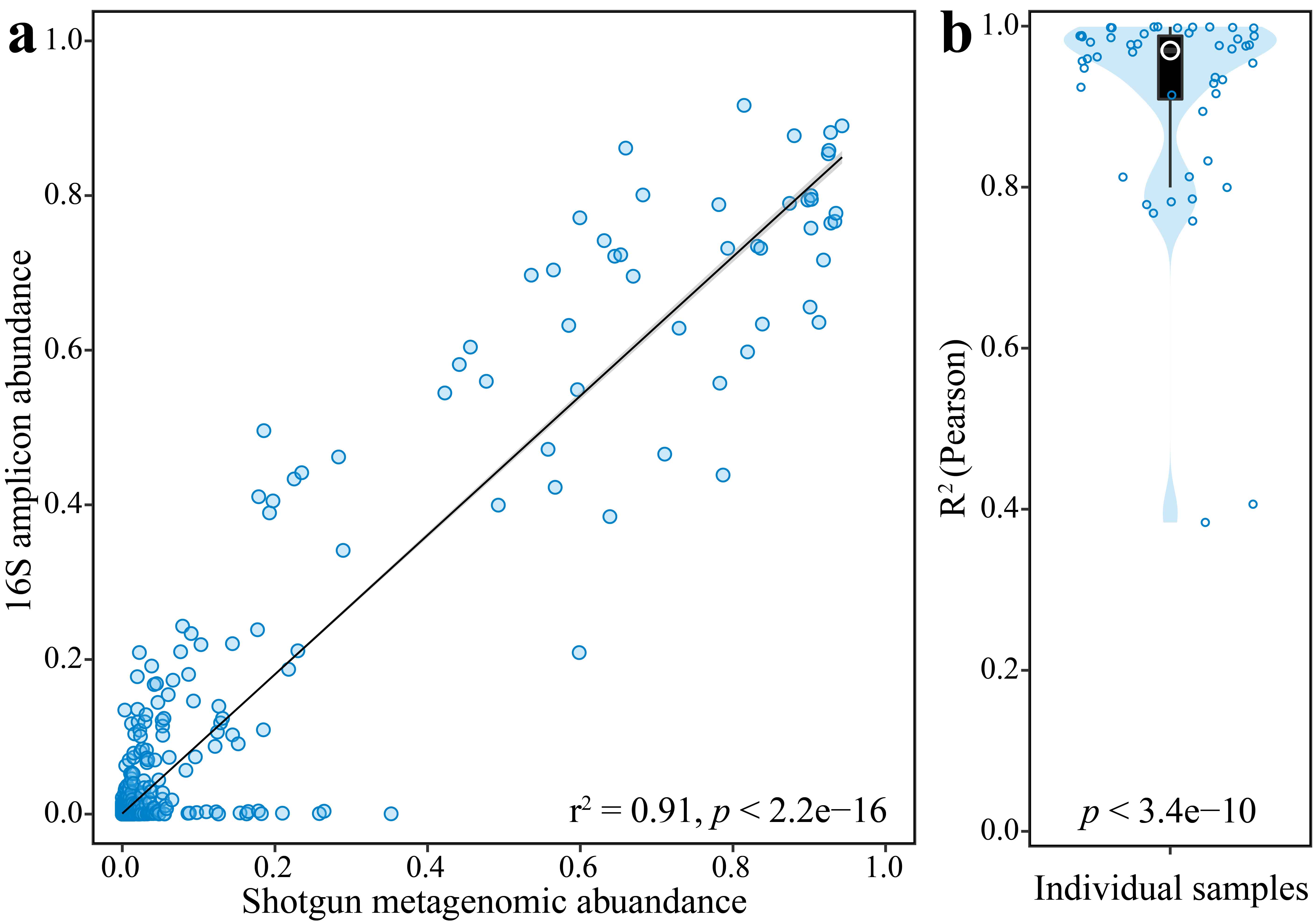

**Fig. S1** Comparison of shotgun metagenome sequencing and 16S amplicon sequencing of genus-level taxa: a) across all investigated samples; b) within individual samples. The 84 genera included in the scatterplot are present in at least one of the enriched samples for both shotgun metagenomic and 16S amplicon sequencing profiles. Line shows the result from simple linear regression (lm function in R), with 95% confidence intervals shown in the shaded areas (plot a). Pearson correlation was applied to calculate the r^2^ value and p-value.

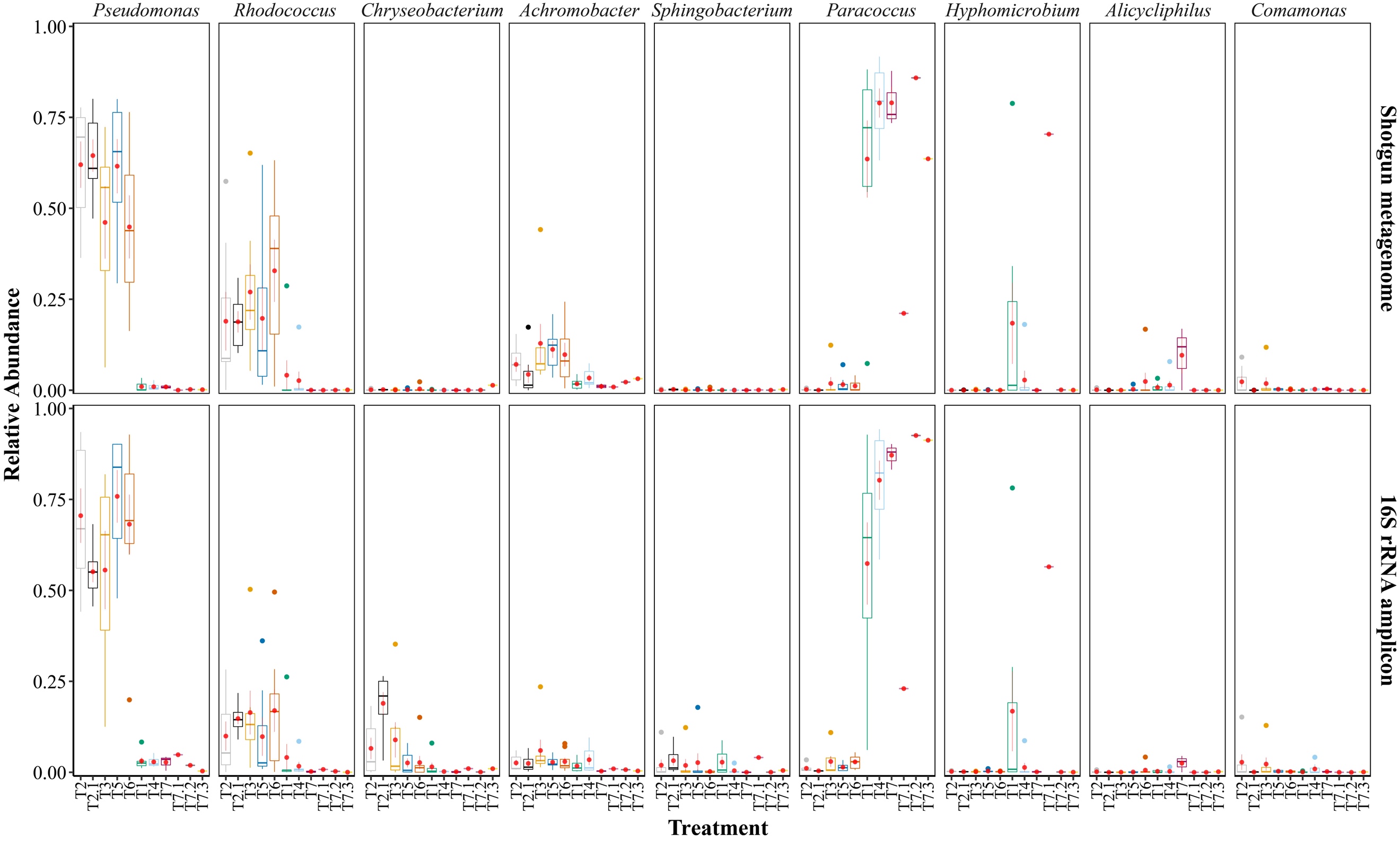

**Fig. S2** Comparative dominant genera revealed by amplicon and metagenomic sequencing approaches. Amplicon results (except T7.1, T7.2, and T7.3 in Gas Station, and *Alicycliphilus* and *Comamonas* in all treatments) were reported previously (Li et al. 2020); T6 in Petrochemical Complex was excluded due to preparation error in shotgun metagenome. T1: Dioxane (DMF), T2: BTEX (DMF), T2.1: BTEX (Dioxane), T3: BTEX + Dioxane (DMF), T4: Dioxane + TCE + TCA (DMF), T5: BTEX + TCE + TCA (DMF), T6: BTEX + Dioxane + TCE + TCA (DMF), T7: TCE + TCA (DMF), T7.1: TCA (DMF), T7.2: TCE (DMF), T7.3: PCE (DMF).

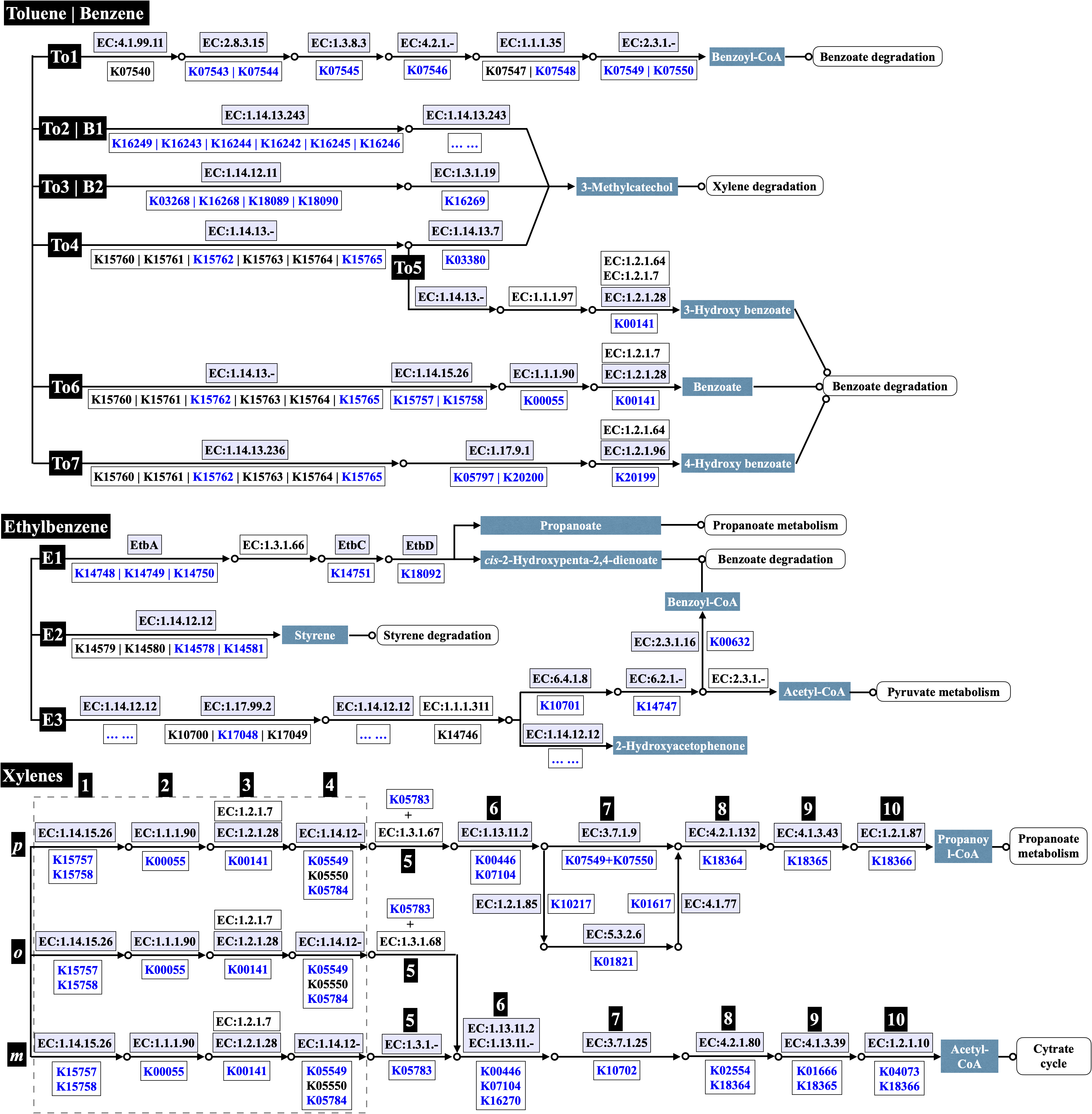

**Fig. S3** Metabolic pathways, enzymes, and associated K numbers involved in the degradation of BTEX. White label with black background indicates contaminant or pathway of contaminant; White label with dark cyan background indicates metabolite produced. K number in blue indicates detected in the microbial communities; K number in black indicates not detected. Enzyme (started with EC) with white background indicates no known associated K number or no associated K number was detected.

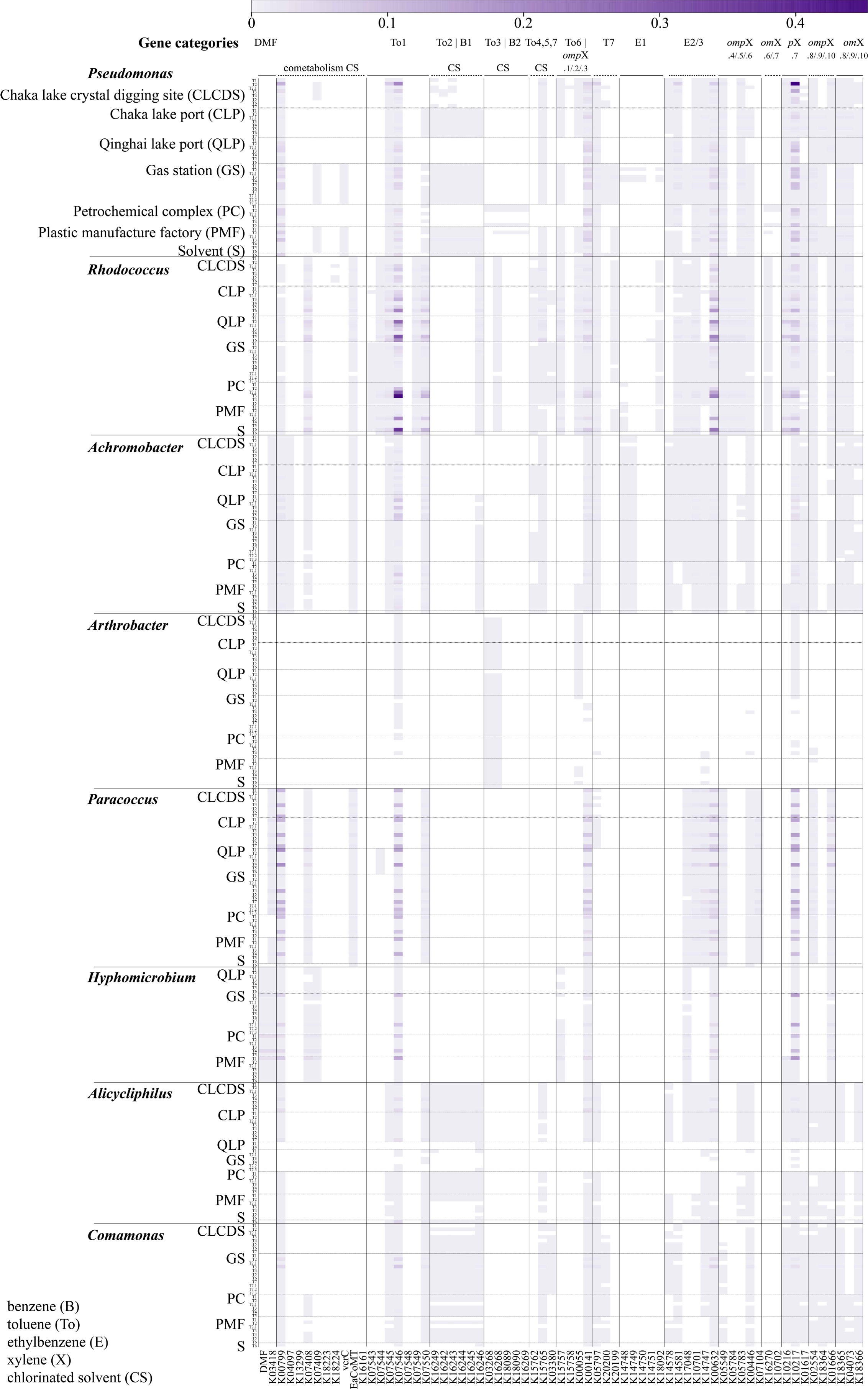

**Fig. S4** Gene abundance of the dominant genera across different treatments. The scale on the top indicates relative gene abundance.

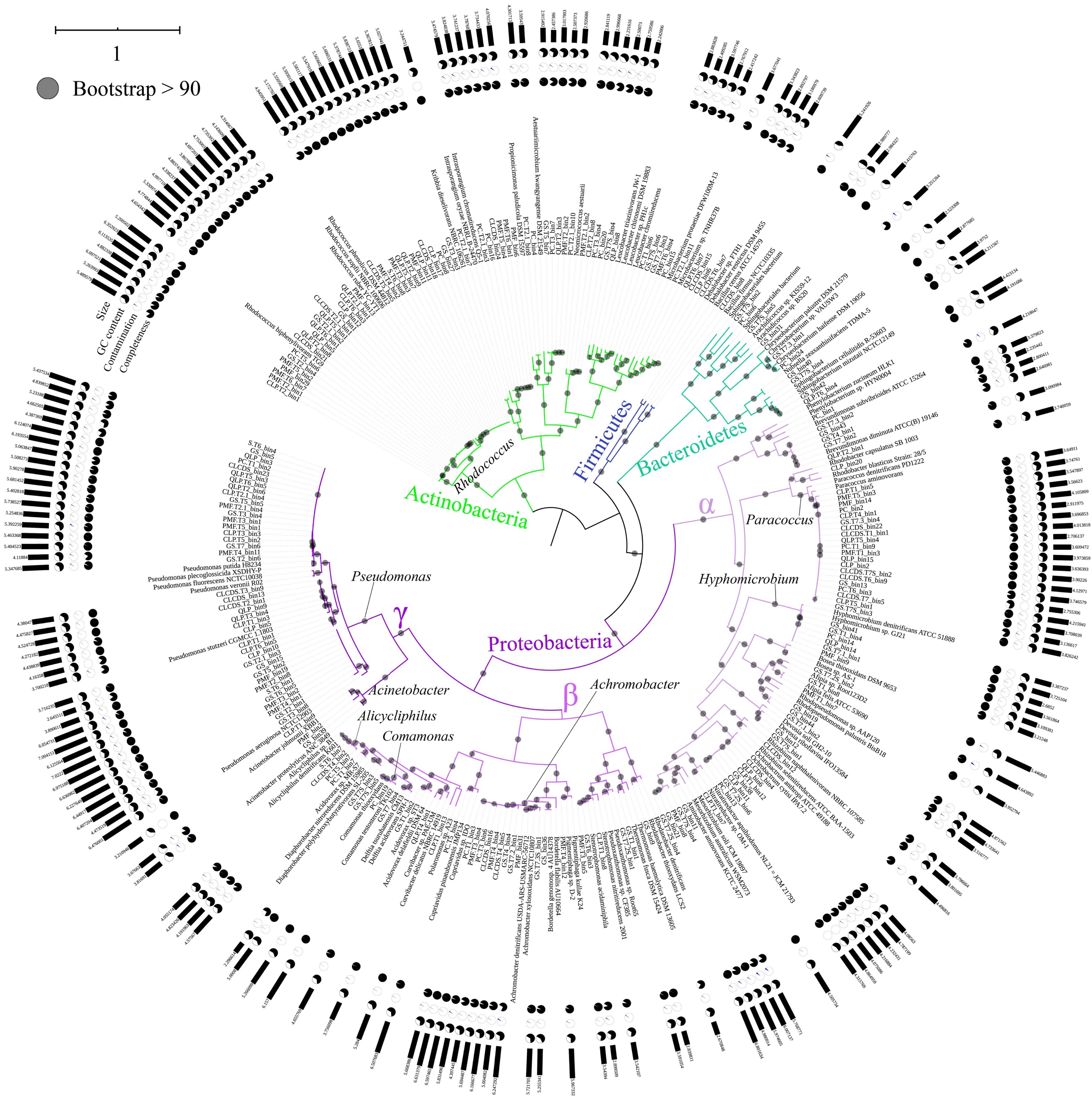

**Fig. S5** Completeness, contamination, GC content, size, and taxonomy of 201 MAGs and 95 GenBank complete genomes with known taxonomy.

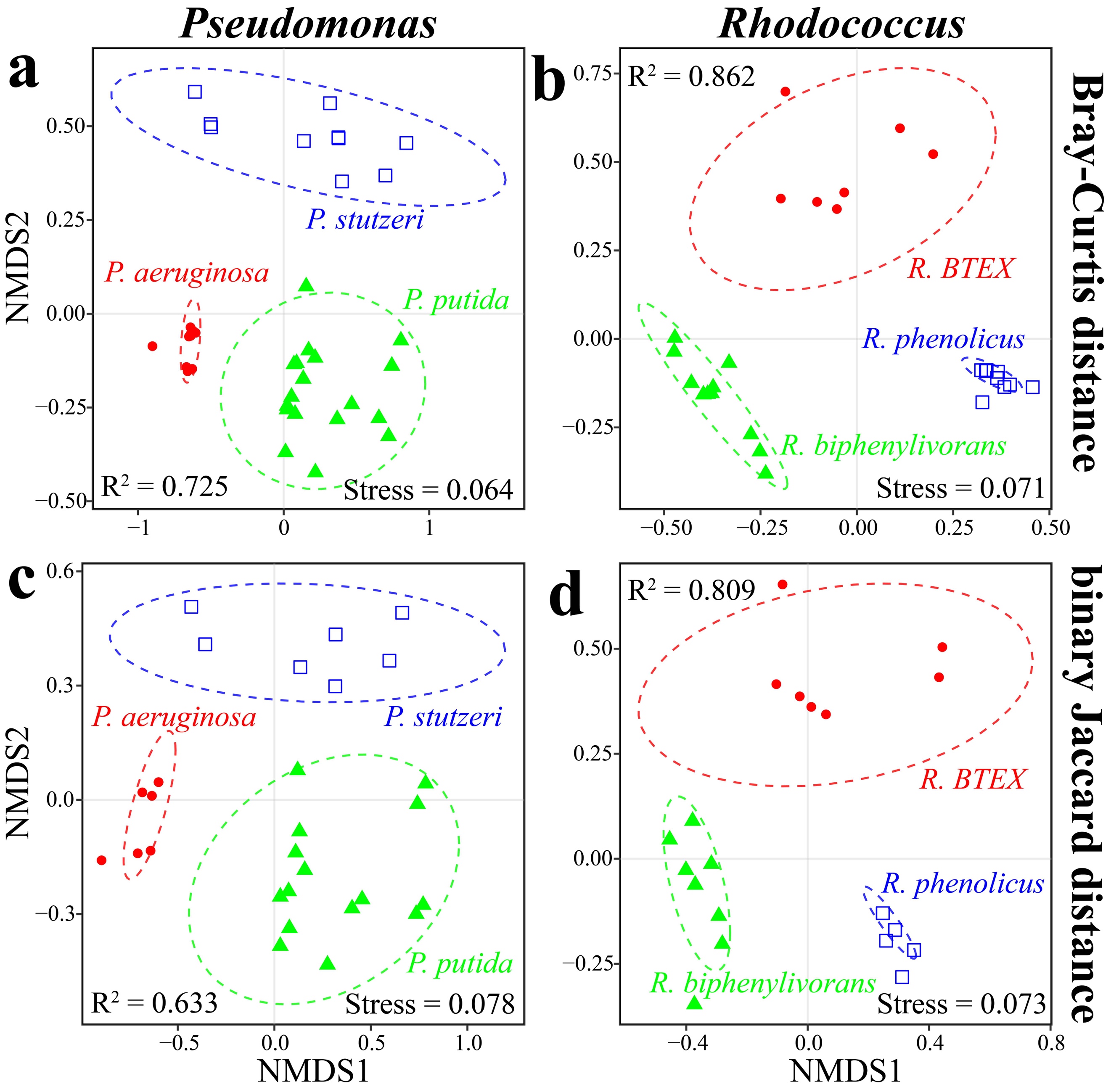

**Fig. S6** Species in the same genus reveal distinct gene repertoires, as indicated by non-metric multidimensional scaling (NMDS) ordination based on Bray-Curtis distance (**a** *Pseudomonas*, **b** *Rhodococcus*) and binary Jaccard distance (**c** *Pseudomonas*, **d** *Rhodococcus*) respectively (permutations = 9,999, *p* < 1e-4). Color indicate different species; dotted line indicates 95% confidence interval.

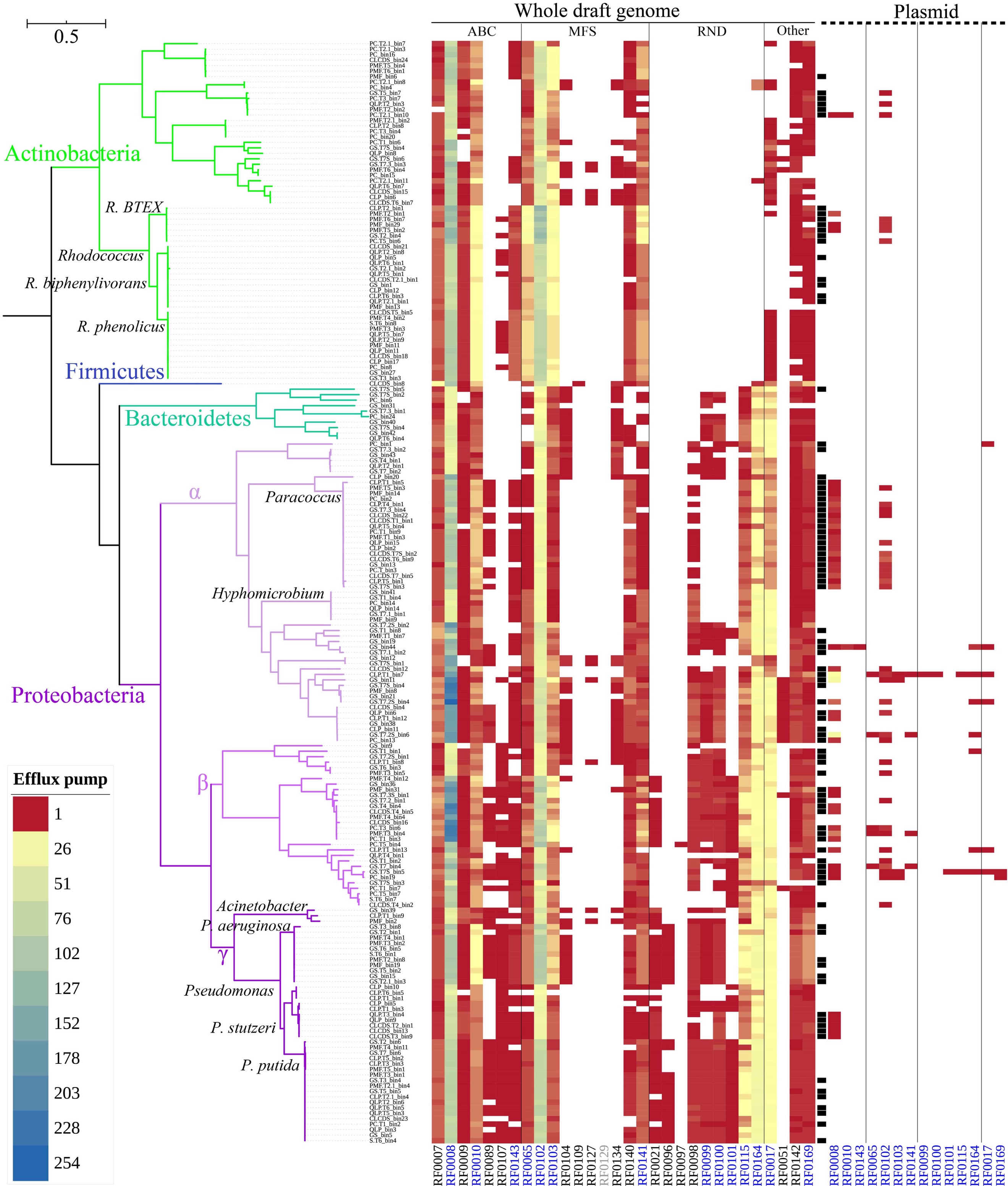

**Fig. S7** Efflux pumps of 201 MAGs. The color gradient denotes the hit of genes associated with efflux pumps in each genome. The column in black in the middle denotes plasmid-containing draft genomes. The RF in blue at the bottom denotes efflux pumps detected in both plasmid and chromosome. RF0129 in grey was not detected in MAGs but was detected in GenBank genomes.

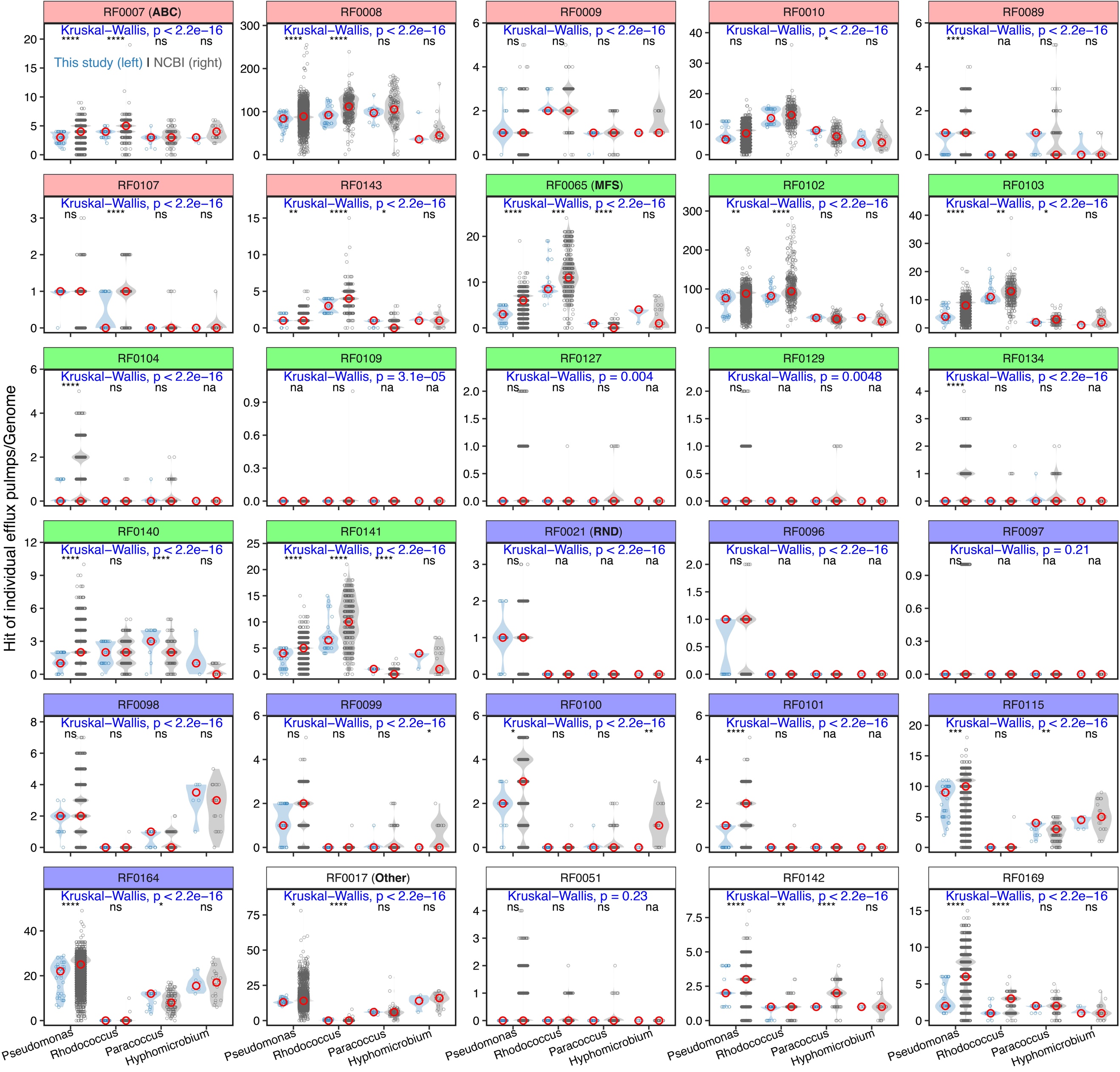

**Fig. S8** Comparison of genome efflux pump genes in four major genera of this study (metagenomic assembled genomes, MAGs) and NCBI (complete and draft) genomes (accessed May 11, 2020). *Pseudomonas* (this study n = 40, NCBI n = 10,352); *Rhodococcus* (this study n = 32, NCBI n = 417); *Paracoccus* (this study n = 20, NCBI n = 132); *Hyphomicrobium* (this study n = 6, NCBI n = 25). Background color of the gene indicates efflux pump category. Red hollow circle indicates median copy number of individual efflux pumps; light blue hollow circle indicates gene copy number of individual efflux pumps of individual genomes in this study; grey hollow circle indicates gene copy number of individual efflux pumps of individual genomes in NCBI. P-value in blue on the plots indicates comparison among four genera, including this study and NCBI (Kruskal-Wallis test). Significance level on the plots indicates comparison between this study and NCBI for each genus (Kruskal-Wallis test): **** indicates *p* < 0.0001; *** indicates *p* < 0.001; ** indicates *p* < 0.01; * indicates *p* < 0.05; ns indicates *p* > 0.05; na indicates zero gene hit.

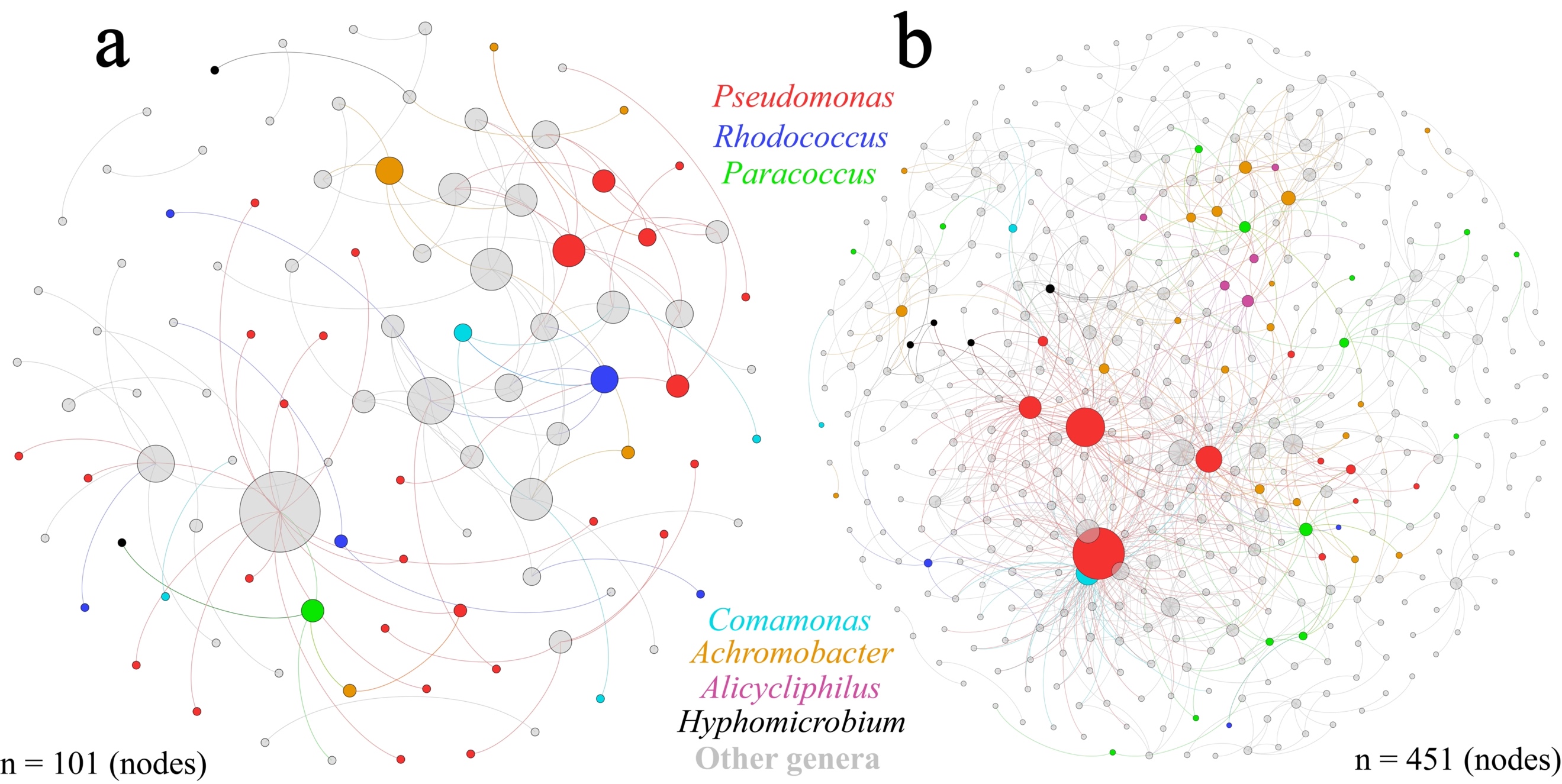

**Fig. S9** Microbial negative co-occurrence network among metagenomic taxa for the treatments in the (**a**) presence and (**b**) absence of BTEX. A connection stands for a strong (Spearman ρ  <  -0.8) and significant (*p* < 0.01, multiple testing adjustment using Benjamini-Hochberg correction) correlation. Color indicates genus. Circle size is proportional to the number of connections (i.e., degree) within each plot.

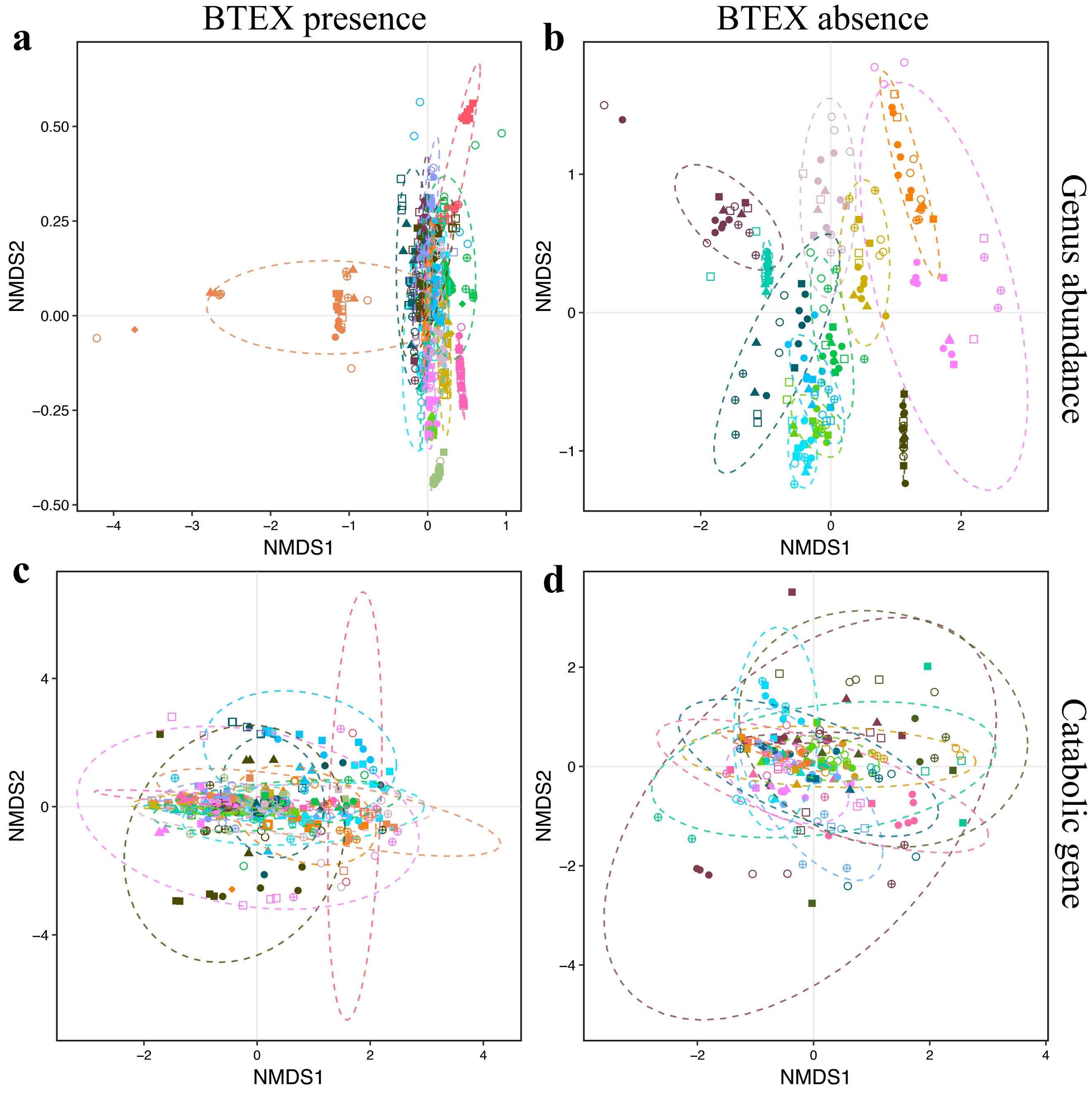

**Fig. S10** Microbial community composition and abundance-weighted catabolic genes differentiate across co-occurrence network modules for the treatments in the presence and absence of BTEX in Fig. 6., as indicated by non-metric multidimensional scaling (NMDS) ordination based on Bray-Curtis distance (permutations = 9,999). (**a**) Genus profiles of microbial community composition in the presence of BTEX, color indicates modules (see Fig. 6a); (**b**) Genus profiles of microbial community composition in the absence of BTEX, color indicates modules (see Fig. 6b); (**c**) Catabolic genes in the presence of BTEX, color indicates modules (see Fig. 6a); (**d**) Genus profiles of in the absence of BTEX, color indicates modules (see Fig. 6b). Shape indicates inoculum source (see Fig. 2). Dotted line indicates 95% confidence interval for individual modules. See Table S4 for Permutational multivariate analysis of variance (PERMANOVA).

**Table S1** Samples used in culturing microbial microbial communities (Li et al. 2020).

| Name | Site | Type | Location | Treatment^#^ | Contaminant exposure history | Incubation period^##^ |
| --- | --- | --- | --- | --- | --- | --- |
| GS | Gas Station | Sub-surface soil | Huizhou, Guangdong, China | T1, T2, T2.1 T3, T4, T5, T6, T7, T7.1, T7.2, T7.3 | Gasoline | Around 16 months |
| PC | Petrochemical Complex | Surface soil | Guangzhou, Guangdong, China | T1, T2, T2.1 T3, T4, T5, T6 | Petroleum hydrocarbons | Around 16 months |
| PMF | Plastic Manufacture Factory | Surface soil | Dongguan, Guangdong, China | T1, T2, T2.1 T3, T4, T5, T6 | Diphenyl carbonate, bisphenol A | Around 16 months |
| S | Solvent contaminated site | Surface soil | Dongguan, Guangdong, China | T6 | Chlorinated solvents, oil | Around 6 months |
| CLCDS | (Hypersaline) Chaka Lake Crystal Digging Site | Sediment | Chaka Lake, Qinghai, China | T1, T2, T2.1 T3, T4, T5, T6, T7 | Potentially diesel, gasoline | Around 16 months |
| CLP | (Hypersaline) Chaka Lake Port | Sediment | Chaka Lake, Qinghai, China | T1, T2, T2.1 T3, T4, T5, T6, T7 | Potentially diesel, gasoline | Around 16 months |
| QLP | (Saline) Qinghai Lake Port | Sediment | Qinghai Lake, Qinghai, China | T1, T2, T2.1 T3, T4, T5, T6 | Potentially diesel, gasoline | Around 16 months |

T1: Dioxane (100 mg/L, DMF); T2: BTEX (400 mg/L, DMF); T2.1: BTEX (400 mg/L, dioxane); T3: BTEX + Dioxane (400 + 100 mg/L, DMF); T4: Dioxane + TCE + TCA (100 + 50 + 50 mg/L, DMF); T5: BTEX + TCE + TCA (400 + 50 + 50 mg/L, DMF); T6: BTEX + Dioxane + TCE + TCA (400 + 100 + 50 + 50 mg/L, DMF); T7: TCE + TCA (50 + 50 mg/L, DMF); T7.1: TCA (50 mg/L, DMF), T7.2: TCE (50 mg/L, DMF), T7.3: PCE (50 mg/L, DMF).

**^#^**T7 (5 mg/L TCE + 5 mg/L TCA, DMF) was absent in some samples due to no microbial growth over 7 days of incubation. **^##^**T2.1: 1 month; T7, T7.1, T7.2, T7.3: 15 months, sub-cultured from T6. In order to get good quality draft genomes associated with the degradation of chlorinated compounds, the diluted cell suspension from the treatments (GS.T7, GS.T7.2, GS.T7.3, CLCDS.T7, and CLP.T7) were spread on mineral salt medium (Li et al. 2020) agar plates, containing the chlorinated compounds of the respective treatments. 12 mixed cultures (GS.T7S1, GS.T7S2, GS.T7S3, GS.T7S4, GS.T7S5, GS.T7S6, GS.T7S7, GS.T7.2S, GS.T7.3S1, GS.T7.3S2, CLCDS.T7S, and CLP.T7S) were picked, extracted DNA, and sequenced.

**Table S2** Comparison of plasmid carriage rate between MAGs and NCBI complete genomes of four genera.

| Genus | MAGs | NCBI complete genomes |
| --- | --- | --- |
| *Pseudomonas* | 45.0% (18/40) | 17.7% (110/623) |
| *Rhodococcus* | 37.5% (12/32) | 66.7% (36/54) |
| *Paracoccus* | 100% (20/20) | 90.0% (18/20) |
| *Hyphomicrobium* | 0% (0/6) | 0% (0/4) |

In A% (B/C), A% indicates percentage of plasmid carriage, B indicates genomes carrying plasmid, C indicates total genomes.

**Table S3** Distribution of nodes across modules in the positive microbial co-occurrence network among metagenomic taxa for the treatments in the a) presence and b) absence of BTEX in Fig. 6.

**a: BTEX presence**

| Module | Node | Percentage |
| --- | --- | --- |
| M1 | 86 | 14.5% |
| M2 | 54 | 9.1% |
| M3 | 52 | 8.8% |
| M4 | 48 | 8.1% |
| M5 | 44 | 7.4% |
| M6 | 37 | 6.3% |
| M7 | 30 | 5.1% |
| M8 | 30 | 5.1% |
| M9 | 24 | 4.1% |
| M10 | 20 | 3.4% |
| M11 | 19 | 3.2% |
| M12 | 15 | 2.5% |
| M13 | 15 | 2.5% |
| M14 | 13 | 2.2% |
| M15 | 10 | 1.7% |
| M16 | 9 | 1.5% |
| M17 | 8 | 1.4% |
| Other | 78 | 13.2% |

Other indicates the sum of all other minor modules with < 1% of node

**b: BTEX absence**

| Module | Node | Percentage |
| --- | --- | --- |
| M1 | 108 | 12.2% |
| M2 | 102 | 11.5% |
| M3 | 101 | 11.4% |
| M4 | 92 | 10.4% |
| M5 | 77 | 8.7% |
| M6 | 74 | 8.3% |
| M7 | 73 | 8.2% |
| M8 | 62 | 7.0% |
| M9 | 59 | 6.7% |
| M10 | 46 | 5.2% |
| M11 | 19 | 2.1% |
| M12 | 9 | 1.0% |
| Other | 65 | 7.3% |

**Table S4** Average relative abundance of *Pseudomonas* genus across modules in the positive microbial co-occurrence network among metagenomic taxa for the treatments in the presence of BTEX in Fig. 6c.

| Module | Mean | Standard deviation |
| --- | --- | --- |
| M1 | 1.3% | 5.4% |
| M4 | 0.014% | 0.026% |
| M5 | 36.3% | 14.5% |
| M9 | 5.3% | 9.9% |
| M14 | 16.6% | 10.8% |

**Table S5** Permutational multivariate analysis of variance (PERMANOVA) of (**a**) microbial community compositions at the genus level and (**b**) abundance-weighted catabolic genes based on Bray-Curtis distance (permutations = 9,999) across modules in the positive microbial co-occurrence network for the treatments in the presence and absence of BTEX in Fig. 6.

| **a**: Genus-level taxonomy | | | | | | | | | | | | | |
| --- | --- | --- | --- | --- | --- | --- | --- | --- | --- | --- | --- | --- | --- |
|  | BTEX presence | | | | | |  | BTEX absence | | | | | |
|  | Df | SumOfSqs | R^2^ | F | Pr(>F) | Sig. |  | Df | SumOfSqs | R^2^ | F | Pr(>F) | Sig. |
| Module | 16 | 98.43 | 0.414 | 21.85 | 0.0001 | *** |  | 11 | 40.26 | 0.422 | 13.9 | 0.0001 | *** |
| Treatment | 4 | 1.08 | 0.005 | 0.96 | 0.5877 |  |  | 5 | 1.56 | 0.016 | 1.18 | 0.086 | . |
| Source | 6 | 3.30 | 0.014 | 1.95 | 0.0001 | *** |  | 5 | 2.42 | 0.025 | 1.84 | 0.0001 | *** |
| Residual | 479 | 134.83 | 0.567 |  |  |  |  | 194 | 51.06 | 0.536 |  |  |  |
| Total | 505 | 237.64 | 1 |  |  |  |  | 215 | 95.29 | 1 |  |  |  |
| **b**: Abundance-weighted catabolic gene^#^ | | | | | | | | | | | | | |
|  | BTEX presence | | | | | |  | BTEX absence | | | | | |
|  | Df | SumOfSqs | R^2^ | F | Pr(>F) | Sig. |  | Df | SumOfSqs | R^2^ | F | Pr(>F) | Sig. |
| Module | 16 | 53.03 | 0.339 | 13.96 | 0.0001 | *** |  | 11 | 17.34 | 0.254 | 5.90 | 0.0001 | *** |
| Treatment | 4 | 0.81 | 0.005 | 0.85 | 0.7609 |  |  | 5 | 0.91 | 0.013 | 0.68 | 0.9875 |  |
| Source | 6 | 4.26 | 0.027 | 2.99 | 0.0001 | *** |  | 5 | 2.85 | 0.042 | 2.14 | 0.0002 | *** |
| Residual | 414 | 98.26 | 0.628 |  |  |  |  | 177 | 47.28 | 0.691 |  |  |  |
| Total | 440 | 156.36 | 1 |  |  |  |  | 198 | 68.38 | 1 |  |  |  |

^#^The module of individual samples without any target genes were excluded in PERMANOVA.

**Reference**

Callahan, B.J., McMurdie, P.J., Rosen, M.J., Han, A.W., Johnson, A.J.A. and Holmes, S.P., 2016. DADA2: High-resolution sample inference from Illumina amplicon data. Nature Methods 13, 581.

Gohl, D.M., Vangay, P., Garbe, J., MacLean, A., Hauge, A., Becker, A., Gould, T.J., Clayton, J.B., Johnson, T.J., Hunter, R., Knights, D. and Beckman, K.B., 2016. Systematic improvement of amplicon marker gene methods for increased accuracy in microbiome studies. Nature Biotechnology 34(9), 942-949.

Johnson, J.S., Spakowicz, D.J., Hong, B.-Y., Petersen, L.M., Demkowicz, P., Chen, L., Leopold, S.R., Hanson, B.M., Agresta, H.O., Gerstein, M., Sodergren, E. and Weinstock, G.M., 2019. Evaluation of 16S rRNA gene sequencing for species and strain-level microbiome analysis. Nature Communications 10(1), 5029.

Li, J., Wu, C., Chen, S., Lu, Q., Shim, H., Huang, X., Jia, C. and Wang, S., 2020. Enriching indigenous microbial consortia as a promising strategy for xenobiotics’ cleanup. Journal of Cleaner Production 261, 121234.

Louca, S., Doebeli, M. and Parfrey, L.W., 2018. Correcting for 16S rRNA gene copy numbers in microbiome surveys remains an unsolved problem. Microbiome 6(1), 41.

Prodan, A., Tremaroli, V., Brolin, H., Zwinderman, A.H., Nieuwdorp, M. and Levin, E., 2020. Comparing bioinformatic pipelines for microbial 16S rRNA amplicon sequencing. PLOS ONE 15(1), e0227434.

Rausch, P., Rühlemann, M., Hermes, B.M., Doms, S., Dagan, T., Dierking, K., Domin, H., Fraune, S., von Frieling, J., Hentschel, U., Heinsen, F.-A., Höppner, M., Jahn, M.T., Jaspers, C., Kissoyan, K.A.B., Langfeldt, D., Rehman, A., Reusch, T.B.H., Roeder, T., Schmitz, R.A., Schulenburg, H., Soluch, R., Sommer, F., Stukenbrock, E., Weiland-Bräuer, N., Rosenstiel, P., Franke, A., Bosch, T. and Baines, J.F., 2019. Comparative analysis of amplicon and metagenomic sequencing methods reveals key features in the evolution of animal metaorganisms. Microbiome 7(1), 133.

Yarza, P., Yilmaz, P., Pruesse, E., Glöckner, F.O., Ludwig, W., Schleifer, K.-H., Whitman, W.B., Euzéby, J., Amann, R. and Rosselló-Móra, R., 2014. Uniting the classification of cultured and uncultured bacteria and archaea using 16S rRNA gene sequences. Nature Reviews Microbiology 12(9), 635-645.
